# IRG1/itaconate/NRF2/GSH axis in tumor-associated macrophages drives therapy resistance and immune evasion in BRCA1-deficient breast cancer

**DOI:** 10.1101/2025.10.14.682312

**Authors:** Yabing Nan, Sophie O’keefe, Xiadi He, Qingyu Luo, Xiaowei Wu, Jing Ni, Yutian Zou, Jerryd Marcus Meade, Yutong Li, Renlei Ji, Anwaruddin Mohammad, Pankaj Kumar, Qiwei Wang, Jean J. Zhao

## Abstract

Tumor-associated macrophages (TAMs) are major contributors to immunosuppression and therapeutic resistance, including resistance to PARP inhibitors (PARPi) in BRCA1-deficient breast cancer. However, the mechanisms underlying TAM-mediated PARPi resistance remain unclear. Here, we demonstrate that TAM-derived glutathione (GSH) impairs the efficacy of PARPi by protecting tumor cells from DNA damage and ferroptosis while suppressing STING-mediated immune activation. Mechanistically, STAT5-driven upregulation of the IRG1/itaconate axis in TAMs rewires mitochondrial metabolism and activates NRF2-dependent GSH biosynthesis. GSH is subsequently released into the tumor microenvironment, where it is taken up by tumor cells, protecting them from PARPi-induced cytotoxicity and dampening immune responses. Pharmacological inhibition of IRG1 reverses these effects, restoring PARPi sensitivity and enhancing anti-tumor immunity in BRCA1-deficient tumor models. Collectively, these findings uncover a TAM-specific immunometabolic program that limits PARPi efficacy and highlight the IRG1/NRF2/GSH axis as a promising therapeutic target to improve treatment outcomes in BRCA1-associated breast cancer.

## Introduction

Poly(ADP-ribose) polymerase inhibitors (PARPi) have been approved as adjuvant targeted therapy for patients with breast cancer susceptibility gene 1 and 2 (BRCA1/2)-associated breast cancer^1,2^. While PARPi have demonstrated clinical benefit in patients with early-stage breast cancer, their efficacy in patients with advanced disease remains limited^2–4^, highlighting the urgent need to improve the therapeutic outcomes of PARPi, particularly in more aggressive or treatment-resistant settings. Recent studies, including our own, suggest that the therapeutic efficacy of PARPi relies not only on their induction of synthetic lethality in BRCA-deficient tumor cells but also on their ability to activate and enhance anti-tumor immunity to achieve more profound and durable therapeutic responses^5–13^.

Tumor-associated macrophages (TAMs) constitute a major immune cell population in the tumor microenvironment (TME) of various solid tumors, including BRCA-mutant breast cancers^14–16^. These cells are highly heterogeneous and display remarkable plasticity, adapting their phenotype and function in response to dynamic cues from the evolving TME^17–19^. TAMs often shift toward a pro-tumorigenic, immunosuppressive phenotype as tumors progress, contributing to therapeutic resistance and poor clinical outcomes^20^. Through multiple mechanisms such as promoting angiogenesis, suppressing anti-tumor immunity, and facilitating tumor cell survival, proliferation, and metastasis, TAMs play a pivotal role in tumor progression and drug resistance, including PARPi resistance^14,21–23^.

Given their central role in shaping an immunosuppressive TME, TAMs have emerged as a critical obstacle to effective cancer therapy. Notably, we and others have shown that PARPi-elicited stimulator of interferon genes (STING)-dependent innate and adaptive anti-tumor immune responses are crucial for fully leveraging its anti-tumor potential^5,6,24^. However, accumulating evidence indicates that TAMs can suppress this immunomodulatory function of PARPi, specifically, the activation of STING signaling^6,16,25^. Moreover, TAMs have also been shown to dampen the DNA damage and cytotoxic effects induced by PARPi in tumor cells^6,26^. These findings have provided the rationale for a newly launched clinical trial combining PARP inhibition with macrophage-targeting strategies in patients with BRCA1/2-associated metastatic breast cancer (NCT06488378). While the immunosuppressive role of TAMs in this context is increasingly recognized, the precise mechanisms by which TAMs attenuate the cytotoxic efficacy of PARPi remain poorly understood, underscoring the need for further mechanistic investigation.

In this study, we investigate the mechanisms by which TAMs suppress PARPi-induced tumor cell lethality in BRCA1-deficient breast cancer, aiming to reveal novel therapeutic targets to enhance PARPi efficacy. We find that low-molecular-weight (<3 kDa) components from TAM-conditioned medium (CM) suppress PARPi-induced DNA damage and cell death. Through metabolomic profiling of TAM-CM associated with BRCA1-deficient breast tumors, we demonstrate that TAM-derived glutathione (GSH) plays a key role in mitigating PARPi-induced cytotoxicity by reducing reactive oxygen species (ROS)-mediated DNA damage in tumor cells. We further explore how reduced DNA damage attenuates cyclic GMP-AMP synthase (cGAS)-STING-mediated DNA sensing and downstream anti-tumor immunity during PARPi treatment. In parallel, we delineate the molecular pathways driving metabolic reprogramming in TAMs, focusing on the immune-responsive gene 1 (IRG1), also known as aconitate decarboxylase 1 (ACOD1), and its catalytic product itaconate, which activates nuclear factor erythroid 2-related factor 2 (NRF2) to regulate GSH production. Finally, we evaluate the therapeutic potential of targeting key metabolic pathways to overcome TAM-mediated resistance to PARPi. Collectively, our findings define a previously unrecognized immunometabolic mechanism of PARPi resistance in BRCA1-deficient breast cancer and identify actionable vulnerabilities in TAMs that can be exploited to enhance both tumor cell killing and anti-tumor immunity.

## Results

### TAM-secreted GSH suppresses the response of BRCA1-deficient tumor cells to PARP inhibitors by scavenging reactive oxygen species

The therapeutic efficacy of PARPi relies on the induction of synthetic lethality in tumor cells deficient in homologous recombination (HR) repair, particularly in those harboring BRCA1/2 mutations. Our previous study demonstrated that TAMs confer potent inhibitory effects on PARPi-induced synthetic lethal response in BRCA1-deficient breast cancer, and this protective effect was mediated by macrophage-secreted soluble factors of <3 kDa^6^. In the current study, we aimed to further investigate how TAMs suppress the cytotoxic effect of PARPi. Since cytokines are usually greater than 5 kDa^27^, we reasoned that the bioactive components present in the <3 kDa fractions are more likely to be small-molecule metabolites or peptides. We therefore fractionated the <3 kDa TAM-CM using Sep-Pak C18 cartridges to isolate both the peptide-enriched fractions and metabolite-enriched flow through (Fig. 1a and Extended Data Fig. 1a). Notably, while the peptide-enriched fractions failed to protect BP breast tumor cells (*Brca1^-/-^ Trp53^-/-^*, termed BP), the metabolite-enriched TAM-CM retained significant protective activity against olaparib treatment (Fig. 1a). Thus, we hypothesized that the protective activity of TAMs might be mediated by small-molecule metabolites.

**Fig. 1.**
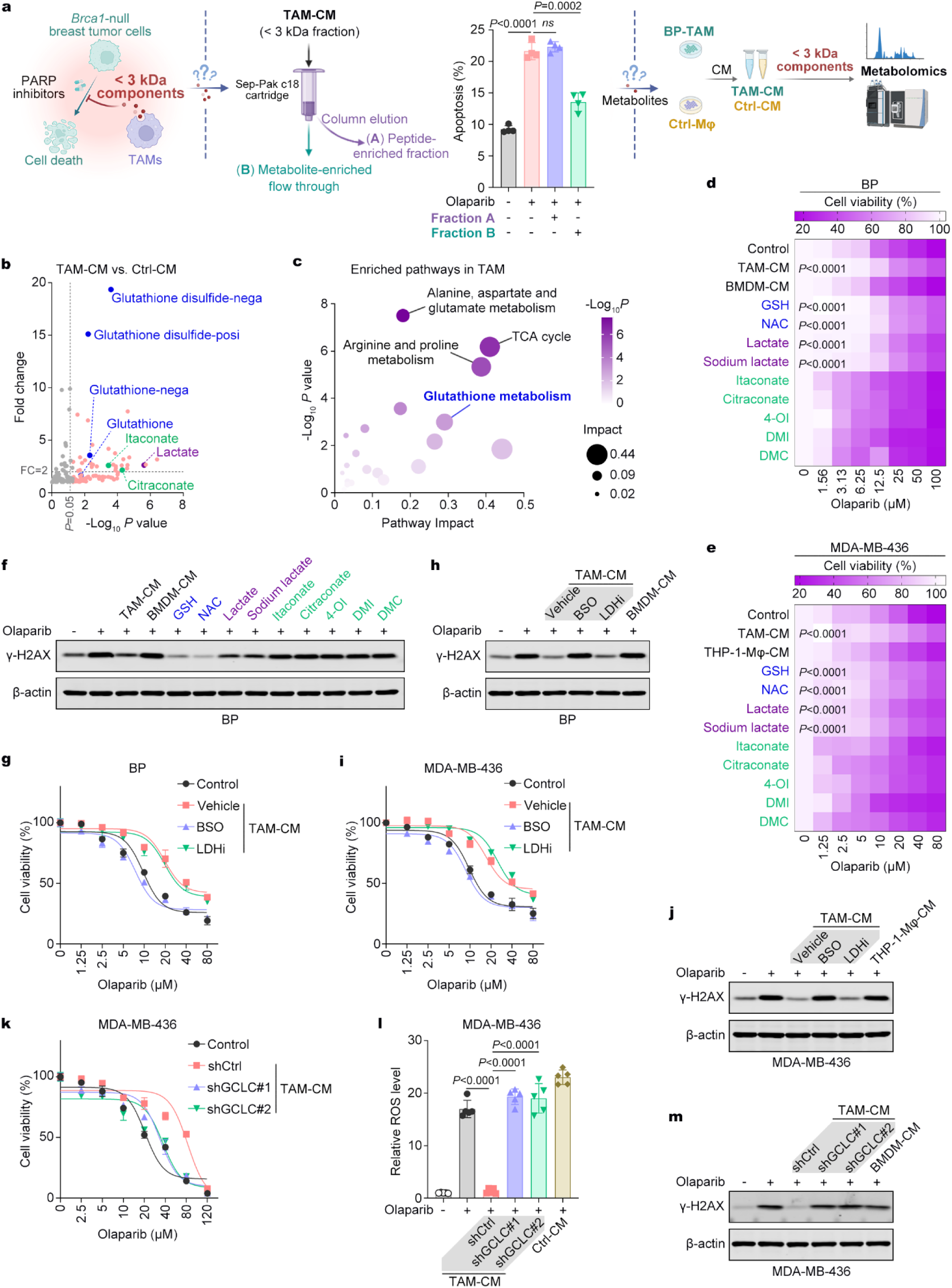
TAM-secreted GSH suppresses the response of BRCA1-deficient tumor cells to PARPi by scavenging ROS. **a**, Schematic illustration of the peptide fractionation strategy from TAM-CM (left); apoptosis levels in BP cells treated with either the peptide-enriched fraction or the metabolite-enriched flow through from TAM-CM in combination with olaparib (middle); and the workflow for untargeted metabolomics of CM derived from control macrophages and TAMs (right). Data are presented as mean ± s.d.; unpaired Student’s t-test; n = 4. **b,** Volcano plots indicate significantly upregulated metabolites in CM derived from BP-TAM compared to control macrophages in the untargeted metabolomic analysis. **c,** Pathway enrichment analysis of metabolites significantly enriched in BP-TAM compared to control macrophages. **d,e,** Heatmap illustrates the relative cell viability of BP cells and MDA-MB-436 cells treated with olaparib at the indicated concentrations for 48 hours, under supplementation with different metabolites. The indicated metabolites, TAM-CM, and BMDM-CM or THP-1 macrophage-CM were added 12 hours before olaparib treatment. The treatment concentrations were GSH and NAC at 5 mM, lactate and sodium lactate at 20 mM, and itaconate and its derivatives at 200 μM. Data represent the mean of three replicates and are color-coded; two-way ANOVA. **f,** Western blot analysis of γ-H2AX expression in BP cells following the indicated treatments. Cells were treated with olaparib (20 μM) for 48 hours, and the metabolites treatment strategy was consistent with (**d**). **g,** Relative cell viability of BP cells treated with olaparib (20 μM) for 48 hours under incubation with different TAM-CM. BMDMs were treated with GSH synthesis inhibitor BSO and lactate synthesis inhibitor sodium oxamate (10 μM) for 48 hours under incubation with BP-CM. CM from the inhibitor-treated TAM and vehicle TAM, as well as control BMDM-CM were used to incubate BP cells for 12 hours before the treatment of olaparib. Data are presented as mean ± s.d.; n = 3. **h,** Western blot analysis of γ-H2AX expression in BP cells treated with olaparib (20 μM) for 48 hours under incubation with different CM. BMDMs were treated with GSH synthesis inhibitor BSO and lactate synthesis inhibitor sodium oxamate (10 μM) for 48 hours under incubation with BP-CM. CM from the inhibitor-treated TAM and vehicle TAM, as well as BMDM-CM were used to incubate BP cells for 12 hours before the addition of olaparib. **i,** Relative cell viability of MDA-MB-436 cells treated with olaparib for 48 hours under incubation with different TAM-CM. THP-1 macrophages were treated with GSH synthesis inhibitor BSO and lactate synthesis inhibitor sodium oxamate (10 μM) for 48 hours under incubation with MDA-MB-436-CM. CM from the inhibitor-treated TAM and vehicle TAM, as well as control THP-1 macrophage-CM were used to incubate MDA-MB-436 cells for 12 hours before the treatment of olaparib. Data are presented as mean ± s.d.; n = 3. **j,** Western blot analysis of γ-H2AX expression in MDA-MB-436 cells treated with olaparib (20 μM) for 48 hours under incubation with different CM. THP-1 macrophages were treated with GSH synthesis inhibitor BSO and lactate synthesis inhibitor sodium oxamate (10 μM) for 48 hours under incubation with MDA-MB-436-CM. CM from the inhibitor-treated TAM and vehicle TAM, as well as control THP-1 macrophage-CM were used to incubate MDA-MB-436 cells for 12 hours before the treatment of olaparib. **k,** Relative cell viability of MDA-MB-436 cells after pre-treated with control or GCLC-knockdown TAM-derived CM for 12 hours, followed by olaparib treatment for 48 hours. Data are presented as mean ± s.d.; n = 3. **l,** Intracellular ROS levels in MDA-MB-436 cells treated with TAM-CM derived from either control or GCLC-knockdown TAMs for 12 hours, followed by co-treatment with olaparib (10 μM) for 48 hours. Data are presented as mean ± s.d.; unpaired Student’s t-test; n = 5. **m,** Western blot analysis of γ-H2AX expression in MDA-MB-436 cells. Cells were pre-treated with TAM-CM derived from either control or GCLC-knockdown TAMs for 12 hours, followed by olaparib (20 μM) treatment for 48 hours before protein collection and analysis.

To investigate this hypothesis, we performed untargeted metabolomics analysis to identify the metabolites enriched in the TAM-CM compared to the CM of control macrophages (Ctrl-CM) (Fig. 1a and Extended Data Fig. 1a). We identified 75 metabolites that were significantly upregulated in TAM-CM (Extended Data Fig. 1b). Among them, GSH-related metabolites exhibited the most significant elevation, along with the increased itaconate and lactate (Fig. 1b). Consistently, pathway enrichment analysis revealed GSH metabolism as one of the significantly activated pathways in TAMs compared to control macrophages (Fig. 1c). To investigate the functions of metabolites enriched in TAM-CM, we treated BRCA1-deficient murine and human breast cancer cells with olaparib in the presence of GSH, N-acetylcysteine (NAC, precursor of GSH), lactate, sodium lactate, or itaconate and its derivatives, including citraconate, 4-octyl itaconate (4-OI), dimethyl-itaconate (DMI), and dimethyl-citraconate (DMC). We found that GSH and NAC, but not itaconate and its derivatives, significantly reduced PARPi-induced synthetic lethality (Fig. 1d,e) and DNA damage response (Fig. 1f and Extended Data Fig. 1c) in both murine and human BRCA1-deficient breast cancer cells, phenocopying the protective effects of TAM-CM. Additionally, while lactate and sodium lactate exhibited some protective effects, they were less potent than GSH and NAC (Fig. 1d-f and Extended Data Fig. 1c). Further analysis of human TAMs similarly revealed elevated GSH levels (Extended Data Fig. 1d,e), indicating that GSH upregulation in TAMs is a conserved feature across species. Furthermore, we blocked the synthesis of GSH or lactate in TAMs using BSO (a GSH synthesis inhibitor) and LDHi (a lactate synthesis inhibitor), respectively. Notably, BSO, but not LDHi, abolished the protective effects of TAM-CM in both murine and human BRCA1-deficient breast cancer cells upon PARPi treatment (Fig. 1g-j), corroborating our findings from metabolite rescue experiments (Fig. 1d-f and Extended Data Fig. 1c). Collectively, our results identify GSH as a key metabolite secreted by TAMs, responsible for suppressing the PARPi-induced lethal response in BRCA1-deficient breast cancer.

Previous studies have shown that reducing oxidative stress contributes to PARPi resistance, highlighting the role of ROS-induced DNA damage in the susceptibility to PARPi^28^. Here, we found that PARPi significantly induced ROS levels and DNA damage in both murine and human BRCA1-deficient breast cancer cells, which was abrogated by exogenous supplementation with GSH or NAC (Extended Data Fig. 1f-i). Given the broad antioxidant capacity of GSH, we reasoned that TAM-derived GSH may function as a ROS scavenger in tumor cells, thus reducing ROS-induced DNA damage and alleviating the reliance on PARP-mediated repair. To test this, we knocked down glutamate-cysteine ligase catalytic subunit (GCLC), the rate-limiting enzyme in GSH synthesis, in macrophages (Extended Data Fig. 1j). Notably, GCLC-silenced TAMs produced significantly less GSH than control TAMs (Extended Data Fig. 1k). Furthermore, CM derived from GCLC-silenced TAMs, rather than control TAMs, failed to inhibit both ROS accumulation and synthetic lethal responses induced by PARPi (Fig. 1k-m). These results suggest that TAM-derived GSH contributes to PARPi resistance by scavenging tumor cell-intrinsic ROS and thereby limiting DNA damage accumulation.

### STAT5 activation drives upregulation of the IRG1/itaconate pathway in TAMs

Given the crucial role of TAM-secreted metabolites, such as GSH, in protecting tumor cells from PARPi treatment, we aimed to investigate the mechanisms underlying the metabolic reprogramming of TAMs. Metabolomic analysis revealed a prominent activation of itaconate metabolism in TAMs, with six itaconate-related metabolites significantly upregulated compared to control macrophages (Fig. 2a,b). The key enzyme driving this metabolic pathway, IRG1, has previously been identified as a pivotal regulator in TAMs, mediating macrophage activation and immunosuppressive functions^29–32^. Indeed, we observed significant upregulation of IRG1 at both the mRNA and protein levels in human BRCA1-deficient MDA-MB-436–associated TAMs (436-TAMs) and mouse BP-TAMs, each relative to their respective control macrophages (Fig. 2c-f).

**Fig. 2.**
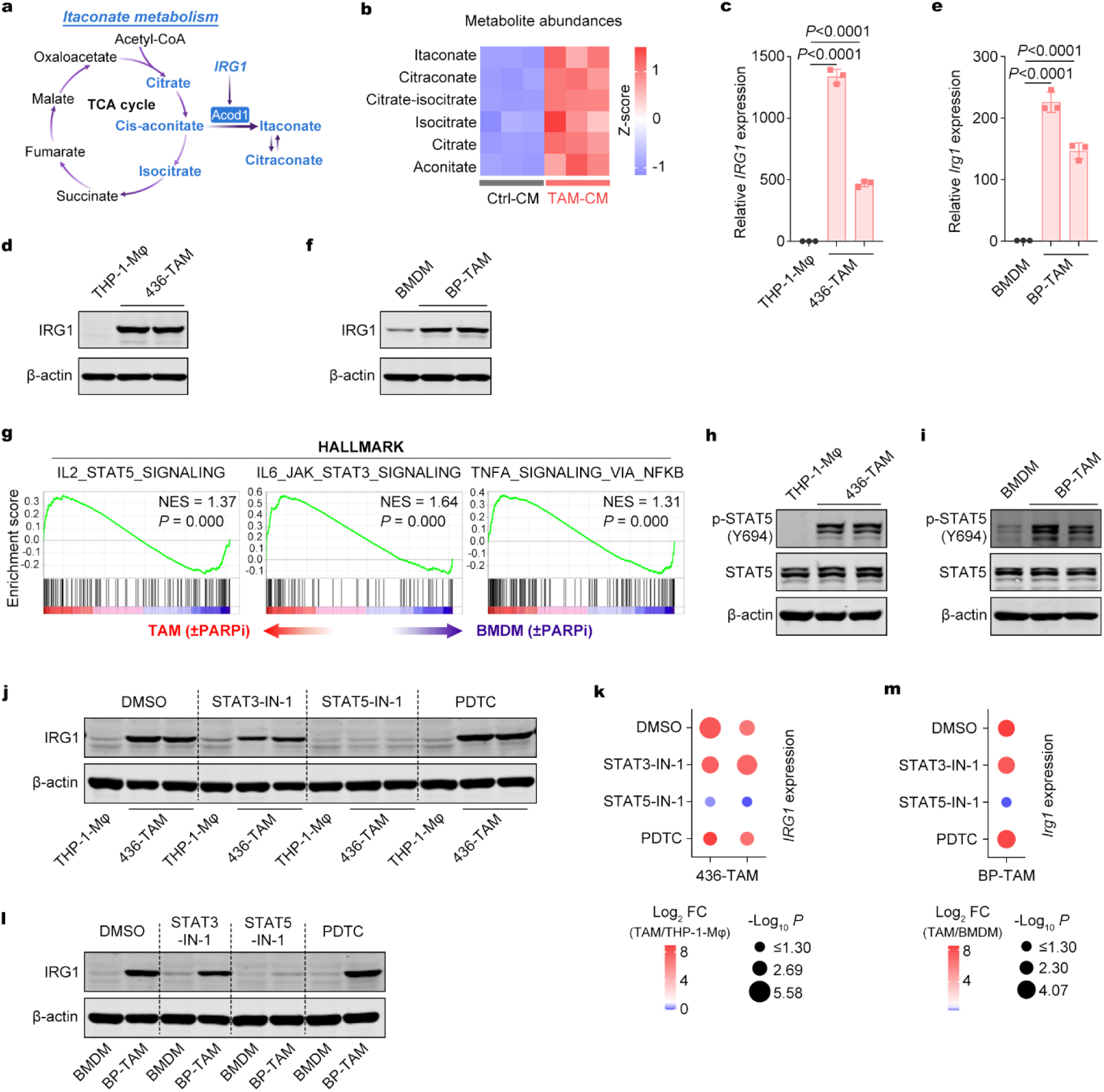
STAT5 activation drives upregulation of the IRG1/itaconate pathway in TAMs. **a**, Schematic representation of the metabolic pathway of IRG1-related metabolites. **b,** Heatmap indicates the relative levels of itaconate and its related metabolites in CM derived from BP-TAM and control macrophages in the untargeted metabolomic analysis. **c,** RT-qPCR analysis of IRG1 expression in THP-1-Mφ and 436-TAMs. Data are presented as mean ± s.d.; unpaired Student’s t-test; n = 3. **d,** Western blot analysis of IRG1 protein expression in THP-1-Mφ and 436-TAMs. **e,** RT-qPCR analysis of IRG1 expression in BMDMs and BP-TAMs. Data are presented as mean ± s.d.; unpaired Student’s t-test; n = 3. **f,** Western blot analysis of IRG1 protein expression in BMDMs and BP-TAMs. **g,** GSEA of RNA-seq data from BMDMs and BP-TAMs. **h,i,** Western blot analysis of STAT5 and phosphorylated STAT5 (p-STAT5) expression in THP-1-Mφ, 436-TAMs, BMDMs, and BP-TAMs. **j,** Western blot analysis of IRG1 protein expression in THP-1-Mφ and 436-TAMs treated with inhibitors of STAT3, STAT5, and NF-κB. The inhibitors were used at the following concentrations: STAT3-IN-1 (10 μM), STAT5-IN-1 (100 μM), and pyrrolidinedithiocarbamate ammonium (PDTC, 10 μM). **k,** RT-qPCR analysis of IRG1 mRNA expression in THP-1-Mφ and 436-TAMs treated with inhibitors of STAT3, STAT5, and NF-κB. Fold changes and *P* values in IRG1 mRNA levels in TAMs relative to THP-1-Mφ are shown. The inhibitors were used at the same concentrations as in (**j**). **l,** Western blot analysis of IRG1 protein expression in BMDMs and BP-TAMs treated with inhibitors of STAT3, STAT5, and NF-κB. The inhibitors were used at the same concentrations as in (**j**). **m,** RT-qPCR analysis of IRG1 mRNA expression in BMDMs and BP-TAMs treated with inhibitors of STAT3, STAT5, and NF-κB. Fold changes and *P* values in IRG1 mRNA levels in TAMs relative to BMDMs are shown. The inhibitors were used at the same concentrations as in (**j**).

We then investigated how IRG1 is upregulated in TAMs. Previous studies suggest that IRG1 can be upregulated in TAMs through the nuclear factor kappa-light-chain-enhancer of activated B cells (NF-κB) pathway^33^ or in tumor-associated neutrophils (TANs) via signal transducer and activator of transcription 5 (STAT5) pathway^34^. Gene set enrichment analysis (GSEA) of our previous RNA-sequencing (RNA-seq) data^6^ revealed significant upregulation of interleukin-2 (IL2)–STAT5, interleukin-6 (IL6)–Janus kinase (JAK)–signal transducer and activator of transcription 3 (STAT3), and NF-κB–tumor necrosis factor alpha (TNFα) signaling in BP-TAMs as compared to control macrophages (Fig. 2g). Western blot analysis further confirmed the significant activation of STAT5, STAT3, and NF-κB in both human and murine TAMs associated with BRCA1-deficient breast cancer (Fig. 2h,i and Extended Data Fig. 2a,b). To determine which pathway(s) drive IRG1 upregulation in TAMs, we employed inhibitors specifically targeting STAT3, STAT5, or NF-κB pathways. STAT5 inhibition completely abolished IRG1 upregulation at both the protein and mRNA levels in human and murine TAMs (Fig. 2j-m). In contrast, the STAT3 inhibitor produced only a modest reduction in IRG1 expression (Fig. 2j,l), and inhibiting NF-κB with PDTC showed no effect (Fig. 2j-m). Together, these findings indicate that STAT5 activation is the primary driver of IRG upregulation in TAMs associated with BRCA1-deficient breast cancer.

### IRG1/itaconate pathway rewires mitochondrial metabolism and promotes GSH synthesis in TAMs

To investigate the impact of IRG1/itaconate pathway activation on TAMs, we examined whether this upregulation induces mitochondrial metabolic reprogramming in these cells. Reanalysis of our transcriptomic data^6^ revealed that, compared to control macrophages, BP-TAMs exhibited significant activation of glycolysis and hypoxia pathways, along with suppressed oxidative phosphorylation (OXPHOS) (Fig. 3a). Consistently, Seahorse assays showed that BP-TAMs exhibited a reduced oxygen consumption rate (OCR) but an elevated extracellular acidification rate (ECAR) compared to control BMDMs (Extended Data Fig. 3a-d). Similarly, 436-TAMs showed markedly suppressed OXPHOS and enhanced glycolysis compared to their control THP1 macrophages (Extended Data Fig. 3e-h). To determine whether this metabolic reprogramming is IRG1-dependent, we generated IRG1-silenced THP-1 macrophages using CRISPR-Cas9 (Fig. 3b). Seahorse assays revealed that the suppressed OXPHOS and enhanced glycolysis observed in 436-TAMs were reversed by silencing IRG1 (Fig. 3c-f), indicating that IRG1 is essential for mediating this mitochondrial metabolic reprogramming in TAMs associated with BRCA1-deficient breast cancer.

**Fig. 3.**
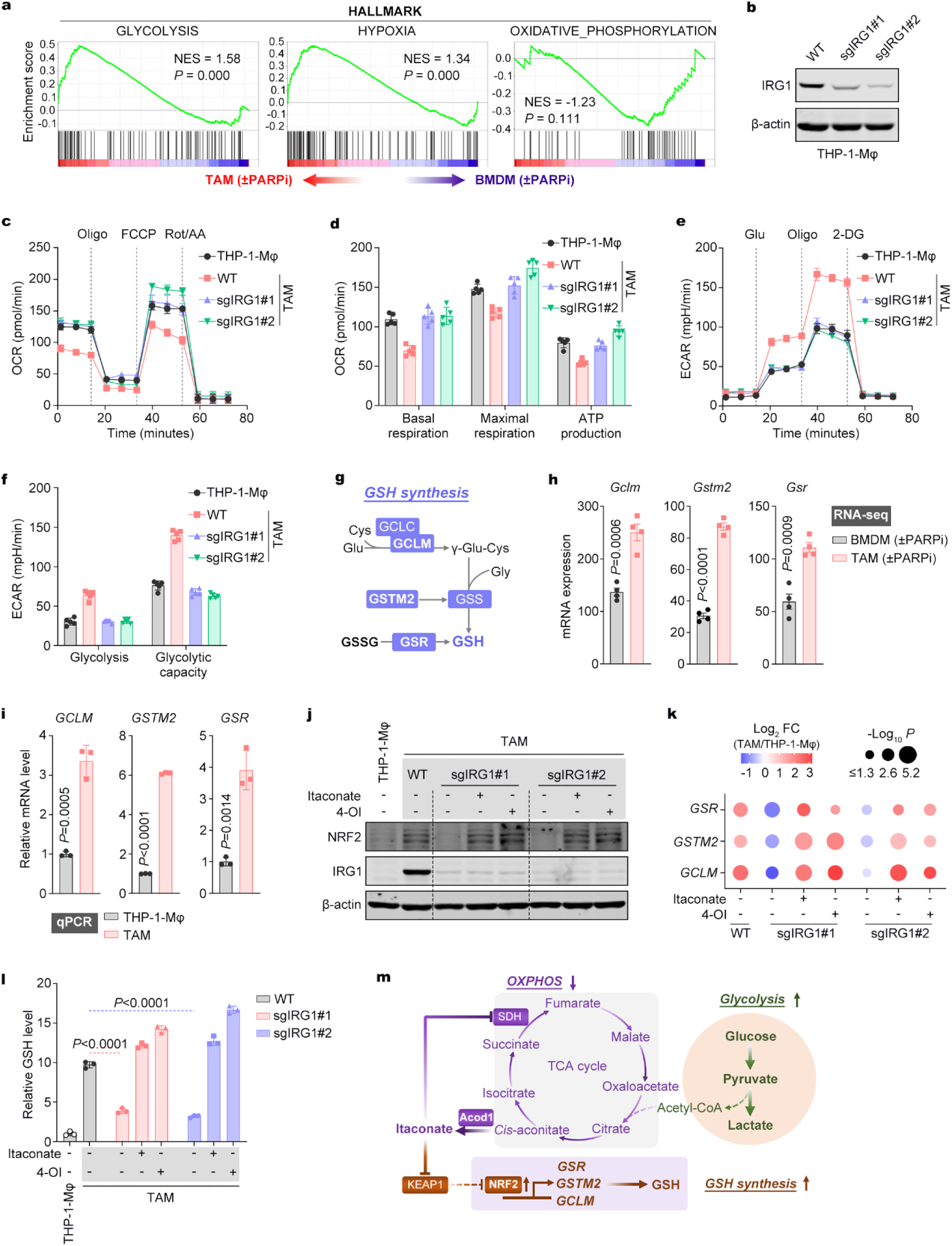
IRG1/Itaconate rewires mitochondrial metabolism and promotes GSH synthesis in TAMs. **a**, GSEA of RNA-seq data from BMDMs and BP-TAMs, with or without olaparib treatment. **b,** Western blot analysis of IRG1 protein expression in THP-1-derived macrophages. **c,d,** Seahorse assays measuring mitochondrial respiration in wild-type and IRG1-knockdown 436-TAMs. Data are presented as mean ± s.d.; n = 5. Oligo, oligomycin; FCCP, carbonyl cyanide *p*-trifluoromethoxyphenylhydrazone; Rot/AA, rotenone and antimycin A; OCR, oxygen consumption rate. **e,f,** Seahorse assays detecting glycolysis in wild-type and IRG1-knockdown 436-TAMs. Data are presented as mean ± s.d.; n = 5. Glu, glucose; Oligo, oligomycin; 2-DG, 2-deoxy-D-glucose; ECAR, extracellular acidification rate. **g,** Schematic illustration of the GSH synthesis pathway and key enzymes involved. **h,** mRNA expression levels of *Gclm*, *Gstm2*, and *Gsr* in BMDMs and BP-TAMs, determined by RNA-seq analysis. Data are presented as mean ± s.e.m.; unpaired Student’s t-test; n = 4. **i,** RT-qPCR analysis of *GCLM*, *GSTM2*, and *GSR* mRNA levels in THP-1-Mφ and 436-TAMs. Data are presented as mean ± s.d.; unpaired Student’s t-test; n = 3. **j,** Western blot analysis of IRG1 and NRF2 protein expression in wild-type and IRG1-knockdown 436-TAMs, with or without itaconate (200 μM) or 4-octyl itaconate (4-OI, 200 μM) treatment for 12 hours. **k,** RT-qPCR analysis of *GCLM*, *GSTM2*, and *GSR* mRNA expression in wild-type and IRG1-knockdown 436-TAMs, with or without itaconate (200 μM) or 4-OI (200 μM) treatment for 12 hours. Fold changes and *P* values of TAMs relative to control macrophages are shown. **l,** Relative GSH levels in wild-type and IRG1-knockdown 436-TAMs, with or without itaconate (200 μM) or 4-OI (200 μM) treatment for 12 hours. Data are presented as mean ± s.d.; unpaired Student’s t-test; n = 3. **m,** Schematic model depicting the IRG1/itaconate-mediated metabolic reprogramming in TAMs. Itaconate suppresses mitochondrial oxidative phosphorylation while enhancing glycolysis. Additionally, itaconate stabilizes NRF2 protein expression, leading to the transcriptional activation of key enzymes involved in GSH synthesis, ultimately promoting GSH production.

Subsequently, we explored whether the IRG1/itaconate pathway also regulates GSH synthesis in TAMs through activating NRF2, a master transcriptional factor of antioxidant responses^35^, whose stability and activity have previously been shown to be enhanced by itaconate^36^. Reanalysis of our previous RNA-seq data^6^ suggested that three key enzymes involved in GSH synthesis, GCLM, GSTM2, and GSR, were significantly upregulated in BP-TAMs compared to control macrophages (Fig. 3g,h). We further validated the significant upregulation of these genes in both human and murine TAMs, compared to their respective control macrophages, using reverse transcription quantitative real-time PCR (RT-qPCR) (Fig. 3i and Extended Data Fig. 3i). To determine whether the upregulation of GSH synthesis-related enzymes in TAMs is mediated by itaconate through the stabilization of NRF2, we examined NRF2 protein levels in TAMs. Western blot analysis showed that 436-TAMs and BP-TAMs significantly increased protein levels of both IRG1 and NRF2 (Extended Data Fig. 3j,k). Importantly, the increases in NRF2 protein, expression of GSH synthesis enzymes, and intracellular GSH levels were observed in wild-type TAMs but not IRG1 knockdown TAMs (Fig. 3j-l). Moreover, supplementation with exogenous itaconate or its derivative 4-OI, both of which are known to stabilize NRF2 protein^36^, restored these effects in IRG1 knockdown TAMs (Fig. 3j-l). In contrast, NRF2 knockdown abrogated the expression of GSH synthesis-related enzymes in TAMs, and these effects could not be rescued by exogenous itaconate or 4-OI (Extended Data Fig. 3l-n), further supporting that NRF2 is the critical downstream effector of itaconate in promoting GSH synthesis. Collectively, these findings demonstrate that itaconate plays a dual role in TAMs: (1) reprogramming mitochondrial metabolism to favor glycolysis, and (2) stabilizing NRF2 to transcriptionally activate the GSH synthesis machinery, thereby enhancing intracellular GSH production (Fig. 3m).

### IRG1/NRF2 pathway-driven GSH production in TAMs inhibits tumor cell ferroptosis elicited by combined GPX4 and PARP inhibition

Recent studies report that glutathione peroxidase 4 (GPX4) inhibitor-induced ferroptosis enhances the sensitivity of tumor cells to PARPi in BRCA1-deficient cancers^37,38^. Interestingly, analysis of drug sensitivity data from the DepMap portal revealed significant positive correlations between the area under the curve (AUC) values of olaparib and multiple well-characterized ferroptosis inducers (erastin, ML210, ML162, and brequinar)^39^ across pan-cancer cell lines (Fig. 4a). Moreover, treatment with RSL3, a GPX4 inhibitor, significantly sensitized MDA-MB-436 and BP cells to PARPi, which was reversed by a potent ferroptosis inhibitor ferrostatin-1 (Fer-1) (Extended Data Fig. 4a,b). These findings confirm a functional interplay between ferroptosis and PARPi-induced cell death in BRCA1-deficient cancer cells.

**Fig. 4.**
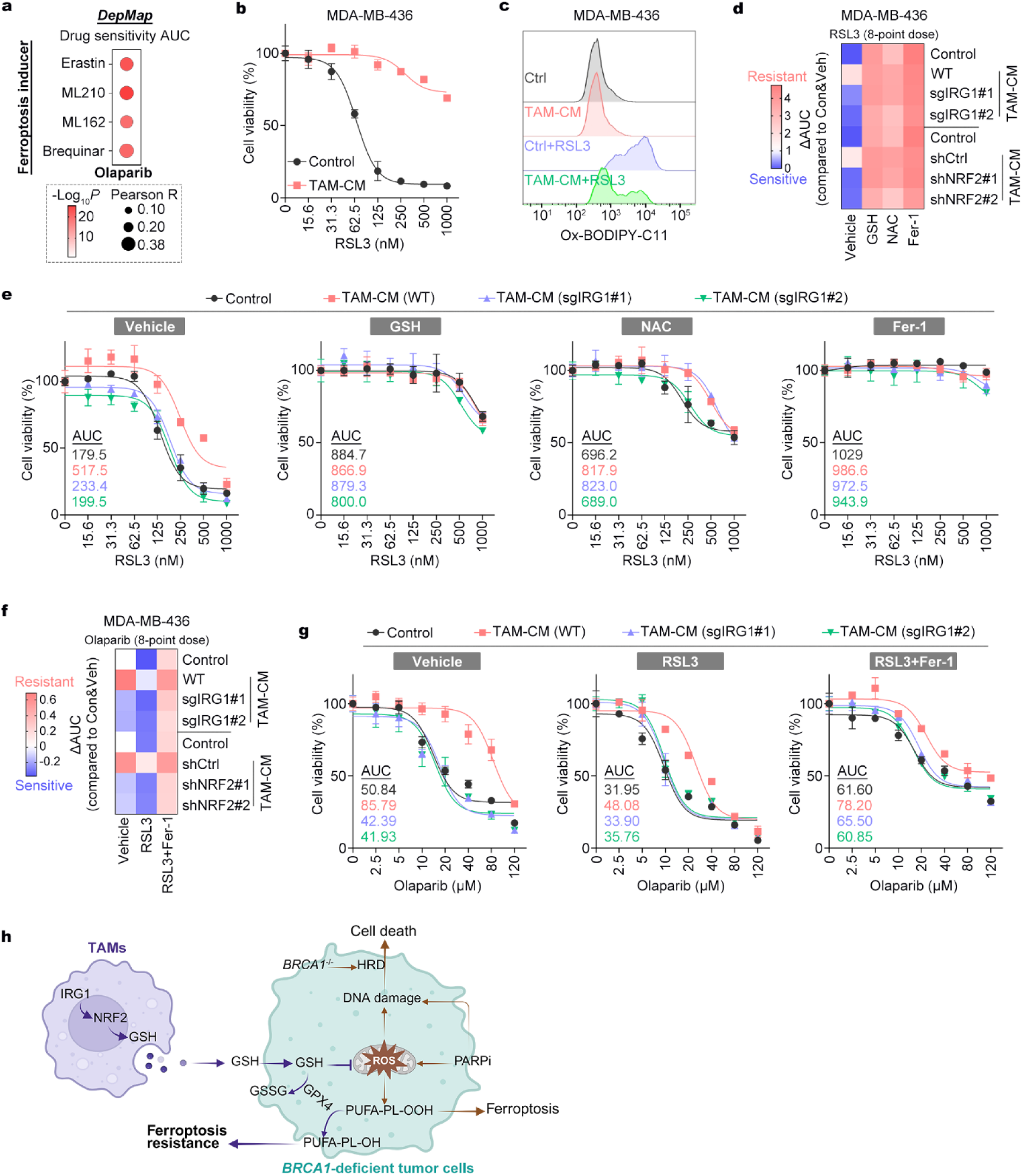
IRG1/NRF2-driven GSH production in TAMs inhibits ferroptosis elicited by combined GPX4 and PARP inhibition. **a**, Correlation analysis of drug sensitivity area under the curve (AUC) values between olaparib and four ferroptosis inducers (erastin, ML210, ML162, and brequinar) across pan-cancer cell lines using the DepMap portal. Pearson correlation coefficients and *P* values are shown. **b,** Drug sensitivity assay assessing RSL3-induced cytotoxicity in MDA-MB-436 cells after pretreatment with control or TAM-CM for 24 hours. RSL3 was then treated at the indicated concentrations for an additional 24 hours before cell viability was measured. Data are presented as mean ± s.d.; n = 3. **c,** Flow cytometry analysis of lipid peroxidation levels in MDA-MB-436 cells pretreated with TAM-CM for 24 hours, followed by co-treatment with RSL3 (250 nM) for 3 hours. **d,** Heatmap depicting ΔAUC values to illustrate the protective effects of GSH (5 mM), NAC (5 mM), and Fer-1 (10 μM) against RSL3-induced ferroptosis. MDA-MB-436 cells were pretreated with TAM-CM from different sources, with or without these compounds for 24 hours, followed by RSL3 treatment for an additional 24 hours before cell viability was assessed. The corresponding viability curves are shown in (**e**) and Extended Data Fig. 4 (**g**). **e,** Dose-response curves and AUC values for MDA-MB-436 cells treated with RSL3 at the indicated concentrations for 24 hours. Treatment conditions were as described in (**d**). Data are presented as mean ± s.d.; n = 3. **f,** Heatmap depicting ΔAUC values to illustrate the effects of RSL3-induced ferroptosis and its inhibition by Fer-1 on olaparib sensitivity. MDA-MB-436 cells were pretreated with TAM-CM from different sources, with or without Fer-1 (10 μM) for 24 hours, followed by co-treatment with RSL3 (100 nM) and olaparib for 48 hours before cell viability was assessed. The corresponding viability curves are shown in (**g**) and Extended Data Fig. 4 (**h**). **g,** Dose-response curves and AUC values for MDA-MB-436 cells treated with olaparib at the indicated concentrations for 48 hours. Treatment conditions were as described in (**f**). Data are presented as mean ± s.d.; n = 3. **h,** Schematic model illustrating the mechanism by which TAMs reduce PARPi sensitivity by conferring ferroptosis resistance in BRCA1-deficient breast cancer. TAM-secreted GSH mitigates PARPi-induced ROS production, thereby reducing DNA damage and lipid peroxidation. Additionally, GSH serves as a substrate for GPX4, further suppressing PARPi/ROS-induced lipid peroxidation, ultimately leading to ferroptosis resistance and PARPi insensitivity.

Given that GSH is a well-established intracellular antioxidant essential for GPX4-mediated ferroptosis defense^39^, we explored whether TAM-derived GSH contributes to ferroptosis resistance in tumor cells. As expected, treatment with GSH or its precursor NAC markedly suppressed RSL3-induced ferroptosis, though their effects were less potent than Fer-1 (Extended Data Fig. 4c,d). Notably, CM from TAMs similarly inhibited RSL3-induced cell death and lipid peroxidation, a hallmark of ferroptosis, in both human and murine BRCA1-deficient breast cancer cells (Fig. 4b,c and Extended Data Fig. 4e,f). To determine whether the IRG1/NRF2 pathway contributes to this protective effect, we analyzed ferroptosis resistance in the context of altered GSH production. CM from wild-type TAMs, but not IRG1-silenced TAMs, protected tumor cells from RSL3-induced ferroptosis (Fig. 4d,e). However, supplementation with GSH, NAC, or the ferroptosis inhibitor Fer-1 largely restored ferroptosis resistance in tumor cells despite IRG1 knockdown (Fig. 4d,e). Similarly, NRF2 knockdown in TAMs abrogated TAM-mediated ferroptosis resistance, which was also reversed by GSH, NAC, or Fer-1 (Fig. 4d and Extended Data Fig. 4g), supporting the role of the IRG1/NRF2/GSH axis in regulating TAM-mediated ferroptosis resistance.

Given that GPX4 inhibitor-induced ferroptosis has been reported to mitigate PARPi resistance in BRCA1-deficient cancers^37^, we next assessed the role of the IRG1/NRF2/GSH axis-mediated ferroptosis resistance in mediating PARPi resistance. Silencing either IRG1 or NRF2 in TAMs abrogated their ability to confer PARPi resistance in BRCA1-deficient tumor cells (Fig. 4f,g and Extended Data Fig. 4h). These findings highlight the importance of the IRG1/NRF2-driven antioxidant program in TAMs, which enhances GSH production and protects tumor cells from PARPi-induced cytotoxicity. Treatment with RSL3 sensitized cancer cells to PARPi and partially attenuated the TAM-mediated protection, although some resistance persisted (Fig. 4f,g and Extended Data Fig. 4h). The addition of Fer-1 restored PARPi resistance even in the presence of CM from IRG1-or NRF2-silenced TAMs (Fig. 4f,g and Extended Data Fig. 4h).

Together, these results highlight the impact of the IRG1/NRF2/GSH pathway in TAMs on protecting BRCA1-deficient breast cancer cells from ROS accumulation and ferroptosis, ultimately enabling resistance to PARPi (Fig. 4h).

### IRG1/NRF2 pathway-driven GSH production in TAMs suppresses PARPi-induced DNA damage and STING activation in tumor cells and dendritic cells

We and others have previously shown that STING-dependent DNA sensing plays a key role in PARPi-mediated anti-tumor immunity^5,8,24^. We further demonstrated that TAMs promote PARPi resistance in BRCA1-deficient breast tumor cells by limiting DNA damage and suppressing STING pathway activation^6^. Consistently, we now show that TAM-derived CM, but not that from control macrophages, markedly suppresses the expression of key STING-dependent cytokine effectors, interferon-beta (IFNβ), C-X-C motif chemokine ligand 10 (CXCL10), and C-C motif chemokine ligand 5 (CCL5)^5,24^, following PARPi treatment in both BP and MDA-MB-436 cells (Extended Data Fig. 5a-d). In addition to tumor cell-intrinsic STING signaling, cytosolic DNA released from damaged tumor cells can also activate the STING pathway in tumor-associated dendritic cells (DCs), a key mechanism for PARPi-induced anti-tumor immunity^5^. Using an *ex vivo* co-culture system, we showed that PARPi-treated tumor cells significantly activated the STING pathway in DCs, as evidenced by concurrently increased phosphorylation of both TANK Binding Kinase 1 (TBK1) and IFN regulatory factor 3 (IRF3), along with upregulation of IFNB, CXCL10, and CCL5 (Extended Data Fig. 5e-g). However, when tumor cells were co-cultured with TAMs, but not with control macrophages, PARPi treatment failed to activate the STING pathway or induce pro-inflammatory cytokines in DCs (Extended Data Fig. 5e-g). These data suggest that TAMs inhibit PARPi-elicited STING pathway activation in both tumor cells and DCs, thereby dampening anti-tumor immune responses.

However, the mechanism by which TAMs suppress PARPi-induced DNA breakage and attenuate STING pathway signaling in tumor cells remains unclear. We therefore investigated whether the IRG1/NRF2/GSH axis in TAMs mediates their inhibitory effect on PARPi-induced STING activation in both tumor cells and DCs. To test this, we collected the CM from TAMs treated with vehicle, IRG1-IN-1 (an inhibitor of IRG1), or ML385 (an inhibitor of NRF2) (Fig. 5a). Western blot analysis revealed that TAM-CM inhibited olaparib-induced TBK1 phosphorylation, whereas control BMDM-CM did not (Fig. 5b). Importantly, this suppression was reversed by inhibition of IRG1 or NRF2, and restored by supplementation with exogenous GSH (Fig. 5b). Consistently, Inhibition of IRG1 or NRF2 abrogated the ability of TAM-CM to suppress *Ifnb*, *Cxcl10,* and *Ccl5* expression as well as IFNβ secretion (Fig. 5c,d). These effects were rescued by exogenous GSH supplementation, reinforcing the role of GSH as a key downstream effector of the IRG1/NRF2 pathway (Fig. 5c,d). We further validated these findings using THP-1-derived TAMs, where inhibition of IRG1 or NRF2 similarly disrupted their ability to suppress STING activation in MDA-MB-436 cells, and this was again restored by exogenous GSH supplementation (Extended Data Fig. 5h-j). Finally, in a tumor cell–DC co-culture system, we confirmed that the IRG1/NRF2 pathway in TAMs governs GSH production, which in turn inhibits PARPi-induced STING activation in DCs (Fig. 5e-g).

**Fig. 5.**
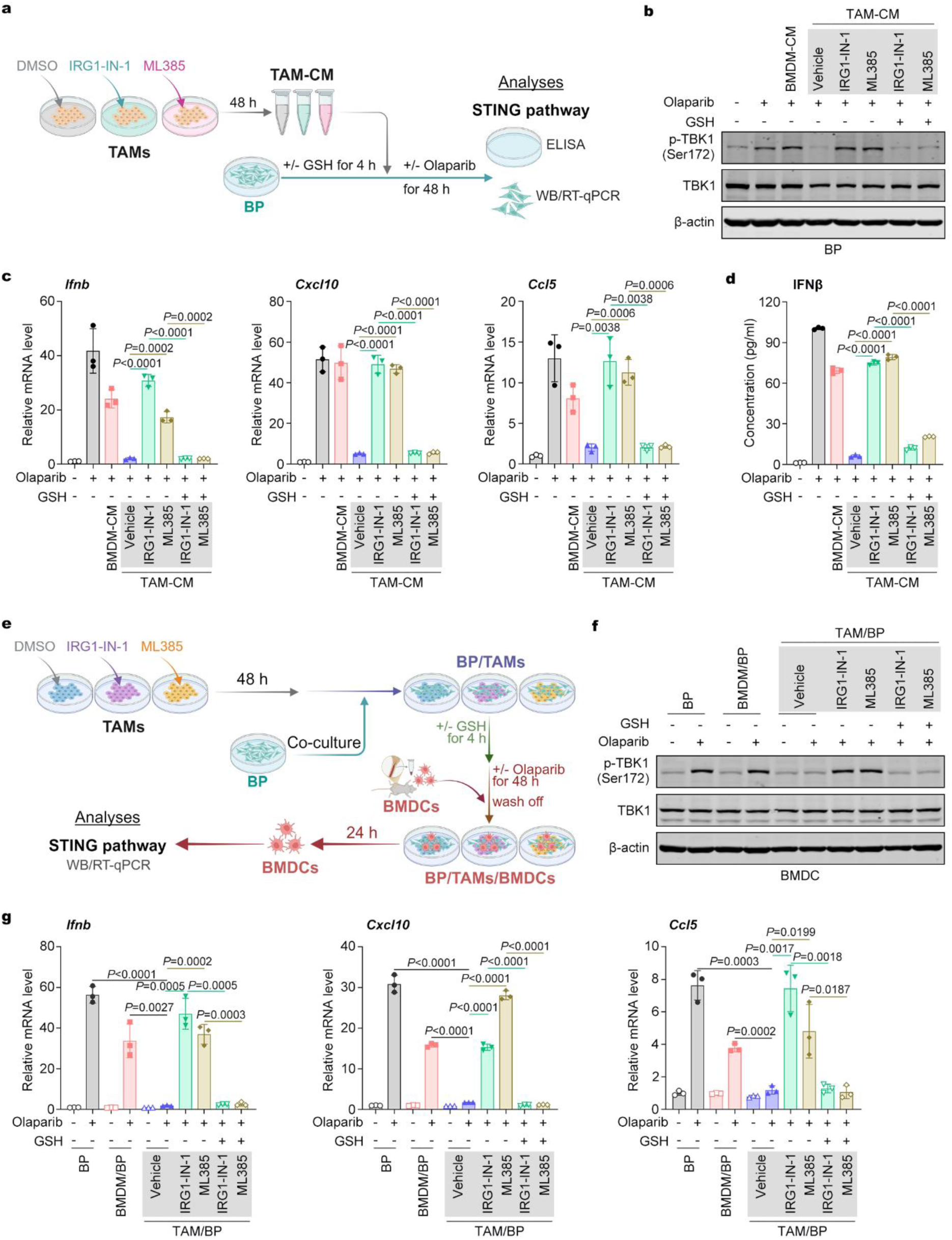
IRG1/NRF2-driven GSH production in TAMs suppresses PARPi-induced STING activation in tumor cells and DCs. **a**, Schematic of the experimental workflow for (**b**) to (**d**). BMDMs were incubated with BP-CM in the presence of DMSO, IRG1-IN-1 (10 μM), or ML385 (10 μM) for 48 hours to generate TAM-CM. BP cells were pretreated with GSH (5 mM) for 4 hours before being treated with different TAM-CM along with olaparib (10 μM) for 48 hours. Supernatants were collected for ELISA, and cells were harvested for western blot and RT-qPCR analysis. **b,** Western blot analysis of TBK1 and p-TBK1 expression in BP cells treated with olaparib and the indicated TAM-CM for 48 hours. GSH was added 4 hours before olaparib treatment. **c,** RT-qPCR analysis of *Ifnb*, *Cxcl10*, and *Ccl5* mRNA levels in BP cells treated with olaparib and the indicated TAM-CM for 48 hours. GSH was added 4 hours before olaparib treatment. Data are presented as mean ± s.d.; unpaired Student’s t-test; n = 3. **d,** ELISA quantification of IFNβ levels in BP cell supernatants following olaparib and TAM-CM treatment for 48 hours. GSH was added 4 hours before olaparib treatment. Data are presented as mean ± s.d.; unpaired Student’s t-test; n = 3. **e,** Schematic of the experimental workflow for (**f**) and (**g**). TAMs treated by DMSO, IRG1-IN-1 (10 μM), or ML385 (10 μM) for 48 hours and then co-cluture with BP cells. This co-culture system was pretreated with GSH (5 mM) for 4 hours, followed by olaparib (10 μM) treatment for 48 hours. Differentiated BMDCs were then added and incubated for an additional 24 hours before being collected for western blot and RT-qPCR analysis. **f,** Western blot analysis of TBK1 and p-TBK1 expression in BMDCs collected after the experimental workflow described in (**e**). **g,** RT-qPCR analysis of *Ifnb*, *Cxcl10*, and *Ccl5* mRNA levels in BMDCs collected after the experimental workflow described in (**e**). Data are presented as mean ± s.d.; unpaired Student’s t-test; n = 3.

Collectively, these findings demonstrate that the IRG1/NRF2-driven GSH production in TAMs suppresses PARPi-induced DNA damage and subsequent STING pathway activation in both adjacent tumor cells and DCs. Targeting this axis in TAMs may enhance STING-mediated immune activation and promote a more immunogenic tumor microenvironment, thereby improving the efficacy of PARPi therapy.

### Targeting IRG1 synergizes with PARPi to suppress BRCA1-deficient breast tumors and reshape the tumor immune microenvironment

To evaluate the *in vivo* therapeutic potential of targeting the IRG1/itaconate/NRF2/GSH axis in TAMs to enhance PARPi efficacy, we tested whether inhibiting this pathway could improve anti-tumor responses in BRCA1-deficient breast cancer. While both classes of IRG1 and NRF2 inhibitors are currently under clinical investigation for cancer, we chose to focus on targeting IRG1, given its more prominent role in TAMs compared to tumor cells, whereas NRF2 functions in both compartments. For this proof-of-concept study, we assessed the combination of IRG1 inhibitor IRG1-IN-1 with olaparib in immunocompetent FVB mice bearing orthotopic syngeneic BP mammary tumors (Fig. 6a). While treatment with either olaparib or IRG1-IN-1 alone had minimal impact on tumor growth, their combination significantly suppressed tumor growth (Fig. 6b). These results were further confirmed in an additional syngeneic model using EO771-sgBRCA1, a BRCA1-knockout breast cancer line generated in our previous studies^6^ (Extended Data Fig. 6a,b). Collectively, these data demonstrate that inhibiting the IRG1-mediated pathway significantly enhances the efficacy of PARPi in BRCA1-deficient breast cancer.

**Fig. 6.**
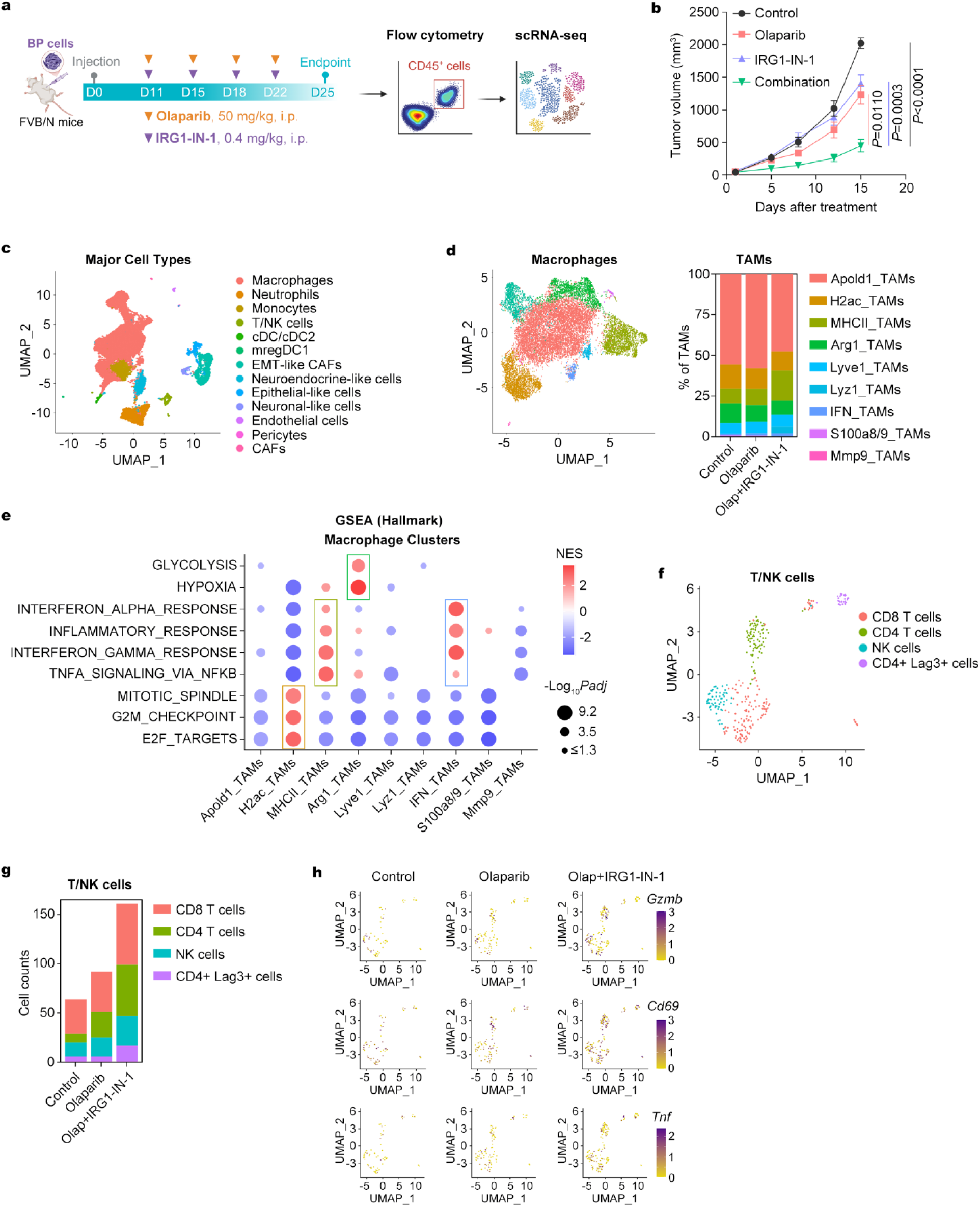
Targeting IRG1 in TAMs enhances the efficacy of PARPi and reshapes TME in BRCA1-deficient breast cancer. **a,b**, Treatment regimen and tumor growth curves of BP tumors. Treatment was terminated when tumors in any group reached 2000 mm^3^. Data are presented as mean ± s.e.m.; two-way ANOVA; n = 8. **c,** Uniform manifold approximation and projection (UMAP) of sorted CD45^+^ cells from tumors treated with vehicle, olaparib alone, or the combination of olaparib and IRG1-IN-1. **d,** UMAP visualization (left) and quantification (right) of TAM subclusters across treatment groups. **e,** GSEA of TAM subclusters. **f,** UMAP visualization of T/NK cell subclusters from tumors under each treatment condition. **g,** Quantification of T/NK cell subclusters from tumors treated with vehicle, olaparib alone, or the combination of olaparib and IRG1-IN-1. **h,** UMAP projection of T/NK cells showing expression of *Gzmb*, *Cd69*, and *Tnf*.

Given our observations that IRG1 inhibition synergized with PARP inhibition to activate STING-interferon signaling in both tumor cells and DCs, we next performed single-cell RNA sequencing (scRNA-seq) on CD45^+^ leukocytes isolated from BP tumors treated with vehicle, olaparib alone, or the combination of olaparib and IRG1-IN-1 (Fig. 6a). Transcriptomic profiles were analyzed using Uniform Manifold Approximation and Projection (UMAP), which revealed 13 major immune cell clusters, with macrophages comprising the dominant population (Fig. 6c), consistent with previous studies showing that advanced BRCA1-deficient breast tumors are heavily infiltrated by pro-tumor TAMs^16^.

To better characterize the macrophage phenotype changes, we performed further sub-clustering analysis and identified nine sub-populations of TAMs based on canonical signatures and differentially expressed genes (Fig. 6d and Extended Data Fig. 6c). Notably, treatment with olaparib alone had minimal impact on the composition of TAMs, with their subtype distribution largely resembling that of the control group (Fig. 6d). In contrast, combining olaparib with the IRG1 inhibitor IRG1-IN-1 led to a marked shift in the TAM landscape (Fig. 6d). Specifically, the proportion of immunosuppressive TAM subsets, such as pro-angiogenic (Apold1_TAMs) and hypoxic (Arg1_TAMs) subtypes, were reduced following the combination therapy (Fig. 6d,e and Extended Data Fig. 6c). At the same time, there was a notable expansion of immune-stimulatory TAMs, including MHC-II_TAMs and IFN_TAMs, compared to both control and olaparib alone treated tumors (Fig. 6d,e and Extended Data Fig. 6c).

Finally, we analyzed T/NK cell populations and identified four major subsets: CD8^+^ T cells, CD4^+^ T cells, NK cells, and CD4^+^ Lag3^+^ regulatory T cells (Fig. 6f and Extended Data Fig. 6d). Quantitative analysis of these lymphoid populations revealed that treatment with olaparib alone primarily increased the proportion of CD4^+^ T cells within the tumor microenvironment (Fig. 6g). In contrast, the combination of olaparib with the IRG1 inhibitor IRG1-IN-1 led to a robust expansion across all lymphoid subsets, including CD8^+^ T cells, CD4^+^ T cells, and NK cells, relative to control or olaparib alone treated tumors (Fig. 6g). Moreover, tumor-infiltrating T cells in combination-treated tumors displayed elevated expression of cytotoxic cytokines (*Gzmb* and *Tnf*) and T cell activation marker (*Cd69*) (Fig. 6h and Extended Data Fig. 6e), suggesting enhanced T cell functionality.

Together, these findings demonstrate that targeting IRG1 not only enhances the cytotoxic effects of PARPi but also reshapes the TME toward an immune-activated, pro-inflammatory state in BRCA1-deficient breast tumors.

## Discussion

Given the mounting evidence that TAMs contribute to resistance against a wide range of anti-cancer therapies, there has been increasing interest in targeting pro-tumorigenic TAMs as a strategy to improve therapeutic outcomes^40,41^. Early efforts have focused on depleting or reprogramming TAMs using monoclonal antibodies or small-molecule inhibitors directed against colony-stimulating factor 1 (CSF1)/CSF1 receptor (CSF1R)^16,42,43^ or transforming growth factor beta (TGFβ)/TGFβ receptor (TGFβR)^44,45^ signaling pathways. While these approaches have shown promise in preclinical models, their translation into clinical benefit has been modest, with many agents failing to produce significant responses in early-phase clinical trials^46–48^. More recently, toll-like receptors 7 and 8 (TLR7/8) agonists are being explored as a novel class of TAM-targeting agents, which have shown some promising results in reprogramming TAMs and activating TME to enhance the effectiveness of other therapies^49,50^, but their clinical development has been hindered by the risk of systemic toxicities, concerns of autoimmune potential and poor pharmacokinetics^51^.

In this study, we present a distinct strategy to modulate TAMs by targeting a metabolic pathway uniquely rewired in TAMs from BRCA1-deficient tumors: the IRG1/itaconate/NRF2/GSH axis. Rather than depleting TAMs, we exploit their metabolic vulnerabilities to impair their pro-tumorigenic function. Building upon our previous findings, we show that TAMs secrete small-molecule metabolites (<3 kDa) that protect BRCA1-deficient tumor cells from the cytotoxic effects of PARP inhibition. Through metabolomic analysis and functional validation, we identify GSH as the critical metabolite mediating this protective effect. Strikingly, exogenous supplementation with GSH alone was sufficient to replicate TAM-induced resistance to PARPi in tumor cells, indicating that GSH is a dominant effector of this resistance phenotype.

Mechanistically, this protective redox buffering is driven by an immunometabolic program within TAMs centered around the IRG1/itaconate/NRF2 pathway. IRG1 catalyzes the production of itaconate, which stabilizes and activates NRF2, a master regulator of antioxidant gene expression. This, in turn, upregulates the synthesis of GSH, which is secreted by TAMs into the TME. Once in the extracellular milieu, GSH is taken up by neighboring tumor cells, where it suppresses ROS accumulation, prevents DNA damage, and thereby dampens cGAS-STING-mediated innate immune signaling.

Our data reveal a dual mechanism of TAM-mediated resistance to PARPi: (1) metabolic protection through GSH-mediated suppression of DNA damage and ferroptosis, and (2) immune evasion through inhibition of STING-driven type I interferon responses in both tumor cells and DCs. Although itaconate itself has been reported to modulate cancer cell behavior^31,32^, we found that its direct impact on BRCA1-deficient breast tumor cells was modest, likely due to the low expression of the itaconate transporter solute carrier family 13 member 3 in these cells. Instead, itaconate’s principal role appears to lie within TAMs, where it serves as a signaling metabolite that activates NRF2 and sustains high levels of GSH production.

Importantly, our findings uncover ferroptosis suppression as a key component of GSH-mediated resistance within the TAM-tumor crosstalk in BRCA1-deficient tumors. Ferroptosis, a regulated form of cell death triggered by lipid peroxidation and GPX4 inactivation^52^, is increasingly recognized for its potential to release immunogenic damage-associated molecular patterns (DAMPs) and promote anti-tumor immunity^53–56^. By exporting GSH to the tumor compartment, TAMs effectively mitigate ferroptosis, thereby supporting tumor cell survival and attenuating immune recognition.

Given the central role of IRG1 in orchestrating this immunosuppressive metabolic program, we tested the therapeutic potential of IRG1 inhibition *in vivo*. Using the IRG1-specific inhibitor IRG1-IN-1, we demonstrated that pharmacologic blockade of IRG1 synergizes with PARPi to suppress tumor growth in multiple immunocompetent models of BRCA1-deficient breast cancer, including genetically engineered mouse models and syngeneic BRCA1-knockout cell line-derived tumors. This combination therapy not only improved tumor control but also reshaped the immune landscape, reducing pro-angiogenic and immunosuppressive TAM subsets while expanding MHC-II^+^ and interferon-responsive macrophages. Additionally, NK cell and T cell compartments shifted toward a more inflammatory, cytotoxic phenotype, with increased expression of granzyme B, TNF, and activation markers.

Together, our study uncovers an immunometabolic mechanism by which TAMs orchestrate resistance to PARPi in BRCA1-deficient breast cancer and highlights the IRG1/NRF2/GSH axis as a tractable target to overcome this resistance (Fig. 7). Therapeutic strategies that inhibit this axis may substantially enhance the efficacy of PARPi and other DNA damage response inhibitors, offering a compelling direction for improving outcomes in patients with aggressive BRCA-mutant breast cancers and potentially other malignancies characterized by macrophage-driven therapy resistance.

**Fig. 7.**
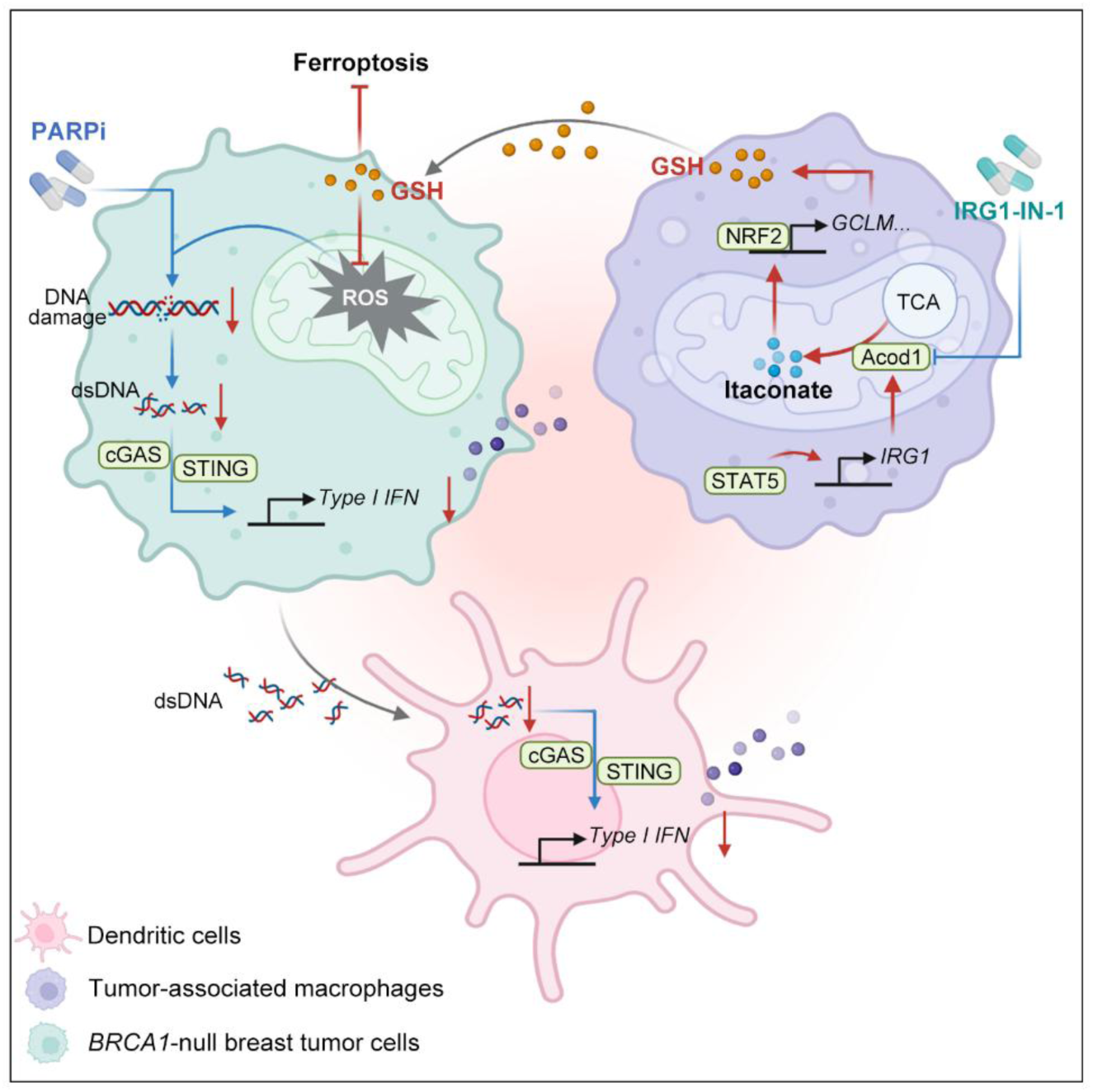
Schematic illustration of the proposed model in which a TAM-mediated immunometabolic axis impairs PARPi efficacy in BRCA1-deficient breast cancer. STAT5-dependent activation of the IRG1/itaconate pathway in TAMs reprograms mitochondrial metabolism and activates NRF2-driven GSH biosynthesis. The TAM-derived GSH protects tumor cells from ROS-induced DNA damage and ferroptosis, while concurrently suppressing STING-mediated immune activation in both tumor and dendritic cells. Pharmacological inhibition of IRG1 restores tumor sensitivity to PARPi, reactivates anti-tumor immunity, and significantly suppresses tumor growth in BRCA1-deficient preclinical models.

## Methods

### Cell culture

The EO771 cell line (940001) was purchased from CH3 BioSystems (Amherst, MA, USA). MDA-MB-436 (HTB-130) and THP-1 (TIB-202) cell lines were obtained from the American Type Culture Collection (ATCC; Manassas, VA, USA). Primary BP tumor cells were isolated from mouse BP mammary tumors reported in our previous study.^6^ EO771 cells were cultured in RPMI 1640 medium (Gibco, Grand Island, NY, USA) supplemented with 10% heat-inactivated fetal bovine serum (FBS; Gemini, West Sacramento, CA, USA) and 10 mM HEPES (15630130, Gibco). MDA-MB-436 cells were maintained in RPMI 1640 medium supplemented with 10% FBS. THP-1 monocytes were cultured in RPMI 1640 medium supplemented with 10% FBS and 0.055 mM 2-mercaptoethanol (21985023, Gibco). All cell cultures were supplemented with 100 μg ml^-1^ penicillin-streptomycin (15140122, Gibco). All cells were maintained at 37 °C in a humidified incubator with 5% CO2. Mycoplasma contamination was routinely tested, and all cell lines were confirmed to be mycoplasma-free. Cell line authentication was performed by short tandem repeat (STR) profiling using the Promega GenePrint 10 System (Promega, Madison, WI, USA).

### Generation of mouse macrophages and dendritic cells

Mouse macrophages and dendritic cells (DCs) were derived from the bone marrow of FVB/N mice and differentiated into bone marrow-derived macrophages (BMDMs) or bone marrow-derived dendritic cells (BMDCs) using distinct induction strategies. Bone marrow cells were seeded into ultra-low attachment plates (Corning, Corning, NY, USA) or standard Petri dishes (Falcon, Corning, NY, USA) and cultured in DMEM supplemented with 10% FBS and 100 μg ml^-1^ penicillin-streptomycin. To induce BMDM differentiation, 10 ng ml^-1^ mouse M-CSF (576404, BioLegend, San Diego, CA, USA) was added to the medium. After three days, the culture was replenished with fresh M-CSF-containing medium, and the cells were incubated for an additional four days. Adherent BMDMs were then harvested for downstream experiments. For BMDC differentiation, bone marrow cells were incubated in DMEM supplemented with 20 ng ml^-1^ mouse GM-CSF (78017, StemCell Technologies, Vancouver, BC, Canada). After three days, the medium was refreshed with GM-CSF, and the cells were further incubated for four more days. Non-adherent BMDCs were subsequently collected for downstream applications.

### Animal experiments

To establish orthotopic mammary tumor models, tumor cells were suspended in serum-free DMEM mixed with 40% Matrigel (Corning) and directly implanted into the mammary fat pads of 6-to 8-week-old female mice. Specifically, female FVB/N mice (001800, The Jackson Laboratory, Bar Harbor, ME, USA) were injected with 5 × 10^5^ BP tumor cells in a total volume of 100 μl. Female C57BL/6J mice (000664, The Jackson Laboratory), received orthotopic injections of 1 × 10^5^ EO771-sgBRCA1 cells in the same volume. Tumor dimensions were regularly measured using digital calipers, recording the longest (length) and shortest (width) diameters. Tumor volumes were estimated using the formula: 0.52 × length × width^2^. Measurements were performed two to three times per week. To minimize inter-operator variability, mice were randomly assigned to treatment groups once the average tumor volume reached approximately 50-100 mm^3^.

Olaparib was administered via intraperitoneal injection at a dose of 50 mg kg^-1^ per day. For each injection, the drug was freshly prepared by diluting a 100 mg ml^-1^ stock solution in DMSO with phosphate-buffered saline (PBS) containing 10% (2-hydroxypropyl)-β-cyclodextrin (HP-β-CD; HY-101103, MedChemExpress, Monmouth Junction, NJ, USA), and injected immediately after preparation. IRG1-IN-1 was administered at a dose of 0.4 mg kg^-1^ per mouse. For each injection, the working solution was prepared by mixing 2 μl of a 5 mg ml^-1^ stock solution with 3 μl DMSO, 30 μl PEG300, 5 μl Tween-80, and 60 μl PBS, resulting in a final injection volume of 100 μl.

### Reagents

The following small-molecule compounds, metabolites, inhibitors, and neutralizing antibodies were used in this study: olaparib (HY-10162, MedChemExpress), L-glutathione reduced (GSH; G4251, Sigma-Aldrich, St. Louis, MO, USA), N-acetyl-L-cysteine (NAC; A7250, Sigma-Aldrich), L-(+)-lactic acid (L1750, Sigma-Aldrich), sodium L-lactate (71718, Sigma-Aldrich), itaconic acid (T4837, TargetMol, Wellesley Hills, MA, USA), citraconic acid (C82604, Sigma-Aldrich), 4-octyl itaconate (4-OI; SML2338, Sigma-Aldrich), dimethyl itaconate (DMI; T5377, TargetMol), dimethyl citraconate (DMC; C0346, TCI America, Portland, OR, USA), DL-buthionine-sulfoximine (BSO; 19176, Sigma-Aldrich), sodium oxamate (LDHi; O2751, Sigma-Aldrich), STAT5-IN-1 (S6784, Selleck Chemicals, Houston, TX, USA), STAT3-IN-1 (S0818, Selleck Chemicals), pyrrolidinedithiocarbamate ammonium (PDTC; S3633, Selleck Chemicals), RSL3 (S8155, Selleck Chemicals), Ferrostatin-1 (Fer-1; S7243, Selleck Chemicals), ML385 (T4360, TargetMol), and IRG1-IN-1 (T78525, TargetMol).

### Virus production and transduction

Lentiviruses were produced in HEK293T cells using the second-generation packaging system with psPAX2 (12260, Addgene, Watertown, MA, USA) and pMD2.G (12259, Addgene). When HEK293T cells reached approximately 95% confluence, the medium was replaced with fresh DMEM without serum or antibiotics. Transfection was performed using Lipofectamine 2000 Transfection Reagent (11668500, Thermo Fisher Scientific, Waltham, MA, USA) according to the manufacturer’s instructions. After six hours of transfection, the medium was replaced with standard DMEM containing 10% FBS and 1× penicillin-streptomycin. Virus-containing supernatants were collected at 48 and 72 hours post-transfection and filtered through a 0.45 µm filter.

For lentiviral transduction, 3 × 10^5^ adherent tumor cells were seeded per well in 6-well plates. Upon cell attachment, 1 mL of viral supernatant and 1 mL of fresh culture medium were added to each well, along with polybrene (TR-1003, Sigma-Aldrich) at a final concentration of 8 μg ml^-1^. Cells were incubated at 37°C for 12 hours, after which the medium was replaced. A second round of infection was performed the following day. After recovery for 24 hours, cells were selected with 2 μg ml^-1^ puromycin (631306, Takara Bio, Shiga, Japan) for approximately one week. Knockdown efficiency was then assessed before proceeding with further experiments. shRNA constructs targeting NRF2 and GCLC were purchased from Sigma-Aldrich: shNRF2#1 (TRCN0000007555): 5’-GCTCCTACTGTGATGTGAAAT-3’, shNRF2#2 (TRCN0000007558): 5’-CCGGCATTTCACTAAACACAA-3’, shGCLC#1 (TRCN0000048484): 5’-GCCATTGAAGAACAATAACTA-3’, shGCLC#2 (TRCN0000048486): 5’-CCAATTCTGAACTCTTACCTT-3’.

### Generation of IRG1-silenced cells

CRISPR-Cas9 genome editing systems were used to generate IRG1-silenced cells^57^. Guide RNA oligonucleotide sequences targeting human IRG1 were designed as follows: *IRG1* (human) #1 forward: 5’-CACCGCGATTACTGCATTTCGCCA-3’; #1 reverse: 5’-AAACTGGCGAAATGCAGTAATCGC-3’. *IRG1* (human) #2 forward: 5’-CACCGGGGCTTCATGCTCCGAAAC-3’; #2 reverse: 5’-AAACGTTTCGGAGCATGAAGCCCC-3’. Annealed guide oligos were cloned into the CRISPR-Cas9 expression vector px458_GFP. The constructed plasmids were transfected into cells. Twenty-four hours post-transfection, single GFP positive cells were sorted into 96-well plates by flow cytometry. After approximately two weeks of culture, emerging colonies were expanded into 6-well plates. IRG1-silenced clones, along with control clones, were validated by western blotting.

### Preparation of tumor cell and TAM conditioned media

For the preparation of conditioned media from BP or MDA-MB-436 tumor cells, cells were cultured until they reached approximately 60% confluence. The original culture medium was then removed, and the cells were washed twice with PBS. A fresh culture medium was added, and the cells were incubated for 48 hours. The supernatant was then collected by centrifugation to obtain tumor cell-conditioned media. To prepare tumor-associated macrophage conditioned media (TAM-CM), BMDMs or THP-1-derived macrophages were first fully differentiated. Cells were then incubated with a 1:1 mixture of tumor-conditioned media (BP-CM or MDA-MB-436-CM) and fresh culture medium for 48 hours. If inhibitor treatment was involved, the inhibitor was added together with the tumor cell-conditioned media during incubation. After 48 hours, the supernatant was collected by centrifugation to obtain TAM-CM from different treatments.

### Cell viability assays

Cell viability was assessed using the CellTiter-Glo® 2.0 Cell Viability Assay (G9242, Promega) according to the manufacturer’s instructions. Drug treatment strategies and incubation times for each experiment are described in the corresponding figure legends. Dose-response curves and area under the curve (AUC) values were generated using a non-linear regression model in GraphPad Prism 9.

### IFNβ secretion detection

The secretion of IFNβ by BP and MDA-MB-436 cells was measured using the LEGEND MAX™ Mouse IFN-β ELISA Kit (439407, BioLegend) and the LEGEND MAX™ Human IFN-β ELISA Kit (449507, BioLegend), respectively, following the manufacturer’s instructions. In brief, tumor cells were treated with the corresponding TAM-CM along with 10 µM olaparib for 48 hours. After incubation, the culture supernatant was collected and centrifuged at 1,500 × g for 10 minutes at 4°C. The pellet was discarded, and the supernatant was used for ELISA analysis.

### Western blotting

Cells were lysed in RIPA lysis and extraction buffer (89901, Thermo Fisher Scientific) supplemented with protease and phosphatase inhibitors and incubated on ice for 10 minutes. The lysates were briefly sonicated and centrifuged at 14,000 × g at 4°C for 10 minutes. The supernatant was collected, and protein concentration was determined using the Pierce BCA Protein Assay Kit (23227, Thermo Fisher Scientific). Protein samples were denatured by boiling at 99°C for 10 minutes. A total of 15 µg of protein per sample was loaded and separated by SDS-PAGE, followed by transfer onto a PVDF membrane. The membrane was blocked with 5% BSA at room temperature for 1 hour, incubated with primary antibodies overnight at 4°C, and then incubated with fluorescent secondary antibodies at room temperature for 2 hours. Protein bands were visualized using an Odyssey® imaging system (LI-COR Biosciences, Lincoln, NE, USA). The following antibodies were used for Western blotting: anti-γ-H2AX (Ser139) (1:1000; 9718S, Cell Signaling Technology, Danvers, MA, USA), anti-GCLC (1:1000; 48005S, Cell Signaling Technology), anti-human IRG1 (1:1000; 77510S, Cell Signaling Technology), anti-mouse IRG1 (1:1000; 19857S, Cell Signaling Technology), anti-NRF2 (1:1000; 12721S, Cell Signaling Technology), anti-STAT5 (1:1000; 94205S, Cell Signaling Technology), anti-p-STAT5 (Tyr694) (1:1000; 9359S, Cell Signaling Technology), anti-STAT3 (1:1000; 9139T, Cell Signaling Technology), anti-p-STAT3 (Tyr705) (1:1000; 9145T, Cell Signaling Technology), anti-NF-κB p65 (1:1000; 8242S, Cell Signaling Technology), anti-p-NF-κB p65 (Ser536) (1:1000; 3033S, Cell Signaling Technology), anti-TBK1 (1:1000; 3504S, Cell Signaling Technology), anti-p-TBK1 (Ser172) (1:1000; 5483S, Cell Signaling Technology), anti-β-actin (1:3000; A2228, Sigma-Aldrich), anti-mouse IgG (1:2000; RL610-145-002, Rockland Immunochemicals, Limerick, PA, USA) and anti-rabbit IgG (1:2000; 35568, Thermo Fisher Scientific).

### RNA preparation and RT-qPCR

Total RNA was extracted using the RNeasy Plus Mini Kit (74134, Qiagen, Hilden, Germany) following the manufacturer’s instructions. Reverse transcription was performed using the High-Capacity RNA-to-cDNA Kit (4387406, Thermo Fisher Scientific) to generate cDNA. qPCR was conducted using the POWER SYBR GREEN PCR Master Mix (4367659, Thermo Fisher Scientific) according to the manufacturer’s protocol. Relative mRNA expression levels were calculated using the ΔΔCT method. Mouse *Actb* and human *GAPDH* were used as endogenous normalization controls for murine and human samples, respectively. The specific qPCR primers used were as follows: human *IRG1* forward 5’-TGGGGCCTTTTATGCCAACT-3’, human *IRG1* reverse 5’-CTCACCTGTGGCCTGTTGAT-3’; human *GCLM* forward 5’-CATTTACAGCCTTACTGGGAGG-3’, human *GCLM* reverse 5’-ATGCAGTCAAATCTGGTGGCA-3’; human *GSTM2* forward 5’-TGTGCGGGGAATCAGAAAAGG-3’, human *GSTM2* reverse 5’-CTGGGTCATAGCAGAGTTTGG-3’; human *GSR* forward 5’-CACTTGCGTGAATGTTGGATG-3’, human *GSR* reverse 5’-TGGGATCACTCGTGAAGGCT-3’; human *IFNB* forward 5’-GCTTGGATTCCTACAAAGAAGCA-3’, human *IFNB* reverse 5’-ATAGATGGTCAATGCGGCGTC-3’; human *CXCL10* forward 5’-GTGGCATTCAAGGAGTACCTC-3’, human *CXCL10* reverse 5’-TGATGGCCTTCGATTCTGGATT-3’; human *CCL5* forward 5’-CCAGCAGTCGTCTTTGTCAC-3’, human *CCL5* reverse 5’-CTCTGGGTTGGCACACACTT-3’; human *GAPDH* forward 5’-CTCTGCTCCTCCTGTTCGAC-3’, human *GAPDH* reverse 5’-TTAAAAGCAGCCCTGGTGAC-3’; mouse *Irg1* forward 5’-AGTTTTCTGGCCTCGACCTG-3’, mouse *Irg1* reverse 5’-AGAGGGAGGGTGGAATCTCT-3’; mouse *Gclm* forward 5’-AGGAGCTTCGGGACTGTATCC-3’, mouse *Gclm* reverse 5’-GGGACATGGTGCATTCCAAAA-3’; mouse *Gstm2* forward 5’-ACACCCGCATACAGTTGGC-3’, mouse *Gstm2* reverse 5’-TGCTTGCCCAGAAACTCAGAG-3’; mouse *Gsr* forward 5’-GACACCTCTTCCTTCGACTACC-3’, mouse *Gsr* reverse 5’-CACATCCAACATTCACGCAAG-3’; mouse *Ifnb* forward 5’-TCCGAGCAGAGATCTTCAGGAA-3’, mouse *Ifnb* reverse 5’-TGCAACCACCACTCATTCTGAG-3’; mouse *Cxcl10* forward 5’-CCAAGTGCTGCCGTCATTTTC-3’, mouse *Cxcl10* reverse 5’-GGCTCGCAGGGATGATTTCAA-3’; mouse *Ccl5* forward 5’-GCTGCTTTGCCTACCTCTCC-3’, mouse *Ccl5* reverse 5’-TCGAGTGACAAACACGACTGC-3’; mouse *Actb* forward 5’-CGGTTCCGATGCCCTGAGGCTCTT-3’, mouse *Actb* reverse 5’-CGTCACACTTCATGATGGAATTGA-3’.

### Intracellular ROS detection

Intracellular ROS levels were measured using the ROS/Superoxide Detection Assay Kit (Cell-based) (ab139476, Abcam, Cambridge, UK) following the manufacturer’s instructions. In brief, 2,000 BP cells or 4,000 MDA-MB-436 cells were seeded into 96-well black-walled, clear-bottom plates. After 12 hours, once the cells adhered, they were treated with 10 µM olaparib in the presence of different TAM-CM or supplemented with metabolites (GSH, 5 mM; NAC, 5 mM) for 48 hours. Following treatment, the medium was replaced with a fresh culture medium, and ROS detection was immediately performed according to the kit protocol. After ROS probe incubation, fluorescence was measured without removing the detection mix using a fluorescence microplate reader.

### Lipid peroxidation detection

Lipid peroxidation was assessed in MDA-MB-436 and BP cells following TAM-CM treatment for 48 hours. The cells were then treated with 250 nM RSL3 for 3 hours, followed by digestion, neutralization, and centrifugation. The cell pellets were washed once with PBS and then resuspended in PBS containing 2 µM BODIPY™ 581/591 C11 (D3861, Thermo Fisher Scientific). The cells were incubated at 37°C for 30 minutes, then washed twice with PBS, and finally resuspended in 200 µL PBS for flow cytometry analysis. Flow cytometry was performed using the FL1 channel, with 10,000 cells analyzed per sample. Data were processed using FlowJo software (BD Biosciences).

### GSH detection

Intracellular and conditioned medium total GSH levels were measured using either the Intracellular Glutathione (GSH) Detection Assay Kit (ab112132, Abcam) or the Glutathione (GSH) Colorimetric Detection Kit (K006-H1, Arbor Assays, Ann Arbor, MI, USA) according to the manufacturer’s instructions. For specific detection of intracellular GSH in tumor cells, cells were incubated with the Thiol Green Dye-loading solution for 30 minutes according to the kit’s protocol, and fluorescence intensity was analyzed using flow cytometry via the FL1 channel. To measure GSH levels in tumor cell-conditioned medium, TAM-CM, and the corresponding TAMs, cells were cultured for 48 hours before the conditioned medium was collected. An equal volume of ice-cold 5% 5-sulfosalicylic acid dihydrate (SSA; S2130, Sigma-Aldrich) was added to the collected medium, followed by incubation at 4°C for 10 minutes. Samples were then centrifuged at 14,000 rpm for 10 minutes at 4°C, and the supernatant was collected for GSH quantification. For intracellular GSH detection in the corresponding TAMs, 5 × 10^6^ TAMs were resuspended in 1 ml of ice-cold 5% SSA, vortexed vigorously to ensure complete cell lysis, and incubated at 4°C for 10 minutes. The lysates were then centrifuged at 14,000 rpm for 10 minutes at 4°C, and the supernatant was collected for subsequent analysis.

### Seahorse assays

Mitochondrial oxidative phosphorylation and glycolysis levels in THP-1-derived macrophages and BMDMs were assessed using the Cell Mito Stress Test Kit (103015-100, Agilent Technologies, Santa Clara, CA, USA) and Glycolysis Stress Test Kit (103020-100, Agilent Technologies) on a Seahorse XFe96 Analyzer (Agilent Technologies). For oxidative phosphorylation analysis, Seahorse XF RPMI medium (103576-100, Agilent Technologies) or Seahorse XF DMEM medium (103575-100, Agilent Technologies) was supplemented with 1 mM pyruvate (103578-100, Agilent Technologies), 2 mM glutamine (103579-100, Agilent Technologies), and 10 mM glucose (103577-100, Agilent Technologies), whereas glycolysis analysis utilized 1 mM glutamine (103579-100, Agilent Technologies) in the base medium. In brief, 8,000 THP-1 cells were seeded into Seahorse XF96 Cell Culture Microplates (101085-004, Agilent Technologies) and differentiated using 100 nM PMA (P1585, Sigma-Aldrich) for 48 hours, followed by a 24-hour recovery in PMA-free medium. The cells were then incubated with 50% conditioned medium from MDA-MB-436 cells for an additional 48 hours before metabolic profiling was performed following the standard Seahorse assay protocol. Similarly, mouse bone marrow cells were seeded into Petri dishes and differentiated into BMDMs with 10 ng ml^-1^ mouse M-CSF (576404, BioLegend) for 7 days. After harvesting and counting, 10,000 BMDMs were seeded per well in Seahorse XF96 Cell Culture Microplates and incubated with 50% conditioned medium from BP cells for 48 hours before metabolic phenotyping. Seahorse XF RPMI medium was used for THP-1 macrophages and Seahorse XF DMEM medium was used for BMDMs. For the mitochondrial stress test, oligomycin, FCCP, and rotenone/antimycin A were sequentially injected at final concentrations of 1.5 μM, 1.0 μM, and 0.5 μM, respectively. For the glycolysis stress test, glucose, oligomycin, and 2-deoxy-D-glucose were sequentially injected at final concentrations of 10 mM, 1 μM, and 50 mM, respectively. Each experimental group included five technical replicates, and data analysis was conducted using Wave software (Agilent Technologies).

### Flow cytometry

Tumors were excised, minced, and enzymatically digested in DMEM supplemented with 5% FBS, 10 mM HEPES, 100 μg ml^-1^ penicillin–streptomycin, 20 μg ml^-1^ DNase I (07900, StemCell Technologies), and 1× collagenase/hyaluronidase (07912, StemCell Technologies) at 37°C for 45 minutes with agitation. Afterward, red blood cells were lysed using ammonium-chloride-potassium (ACK) buffer (BP10-548E, Lonza, Basel, Switzerland), and cell suspensions were filtered through a 70 μm strainer to eliminate undigested tissue fragments.

For flow cytometry, single-cell suspensions were washed and stained in cold FACS buffer (PBS with 0.2% BSA and 5 mM EDTA). Viable cells were identified using the LIVE/DEAD Fixable Aqua Dead Cell Stain (L34965, Thermo Fisher Scientific) for 30 minutes on ice. Cells were subsequently stained with phospho-specific antibodies against p-TBK1 (Ser172) (1:50; 5483, Cell Signaling Technology) or p-IRF3 (Ser396) (1:50; 29047, Cell Signaling Technology) for 30 minutes on ice. To detect phosphorylated H2AX (Ser139), cells were fixed by slow addition of ice-cold 70% ethanol while vortexing, followed by incubation at −20 °C for 1 hour. After washing three times with cold staining buffer, 5 μl of anti-p-H2AX antibody (1:20; 613402, BioLegend) was added to 1 × 10^6^ cells in 100 μl of buffer and incubated at 4 °C for 30 minutes. Samples were acquired using an LSR Fortessa HTS flow cytometer (BD Biosciences). Data were recorded using BD FACSDiva software (version 6) and analyzed with FlowJo (version 10.4).

### Metabolomics

BP (*Brca1* and *Trp53* deleted) breast cancer cells were cultured in serum-free medium for 48 hours to generate conditioned medium. BMDMs were then incubated with the tumor-conditioned medium for 48 hours. Components smaller than 3 kDa were isolated from the incubation medium and subjected to untargeted metabolomic analysis. Differential metabolite analysis and pathway enrichment were performed using MetaboAnalyst 6.0.

### Bulk RNA-seq

The transcriptomic data of BMDMs were obtained from our previously published study.^6^ Gene Set Enrichment Analysis (GSEA) was performed using GSEA 4.3.1 software, referencing the MSigDB Hallmark gene sets for analysis.

### Single-cell RNA-seq and data analysis

Tumor tissues collected from mice treated with vehicle, olaparib alone, or the combination of olaparib and IRG1-IN-1 were enzymatically dissociated into single-cell suspensions using a collagenase/hyaluronidase-based digestion buffer. CD45 positive immune cells were then enriched using a CD45 positive selection kit (18945, StemCell Technologies). For each treatment group, single-cell suspensions from four individual tumors were pooled by centrifugation and resuspended in CryoStor CS10 (100-1061, StemCell Technologies) for preservation. Single-cell RNA sequencing was performed by Novogene using the 10x Genomics platform. Bioinformatic analysis of the single-cell data was conducted by the Bioinformatics Core at the University of Virginia. Raw sequencing data have been deposited in the GEO under accession number GSE300991.

### Statistics and reproducibility

Statistical analyses were performed using GraphPad Prism 9.5.1 (GraphPad Software). Tumor growth data were analyzed using two-way ANOVA. For other analyses, an unpaired two-tailed Student’s t-test was used to compare two conditions. A *P*-value of less than 0.05 was considered statistically significant. Data are presented as means ± s.e.m. or means ± s.d., as specified in the corresponding figure legends. Sample sizes (n) are indicated in the respective figure legends.

### Reporting summary

Further information on research design is available in the Nature Portfolio Reporting Summary linked to this article.

## Data availability

All data reported in this paper will be shared by the lead contact upon request. The single-cell RNA-seq data have been deposited in GEO under accession number GSE300991. Any additional information required to reanalyze the data reported in this paper is available from the lead contact upon request.

## Code availability

This paper does not report the original code.

## Acknowledgements

S.O. is supported by the National Cancer Institute (NCI) Cancer Research Training Grant (T32CA009109) and the University of Virginia Provost Fellowship for graduate students. Q.L. is supported by the Claudia Adams Barr Program in Innovative Basic Cancer Research. X.W. is supported by a National Cancer Center fellowship grant. This work was supported in part by grants from the Department of Defense Breast Cancer Research Program Breakthrough Award HT9425-23-1-0026 (Q.W.), American Cancer Society Institutional Research Grant ACS-IRG 000434 (Q.W.), Breast Cancer Research Foundation (J.J.Z.), and National Health Institute (NIH) P50 CA168504 (J.J.Z), P01 CA269021 (J.J.Z.) and CA210057 (J.J.Z). The graphical abstract was created by BioRender.com.

## Author contributions

Y.N., Q.W., and J.J.Z. conceived and designed the study. Y.N. performed the majority of the experiments. S.O., J.M.M., and Q.W. conducted the animal experiments and collected samples for single-cell analysis. X.H. assisted with metabolomics. Q.L. and X.W. provided support for flow cytometry and Seahorse assays. J.N., Y.Z., Y.L., and R.J. assisted with BMDM isolation. A.M. and P.K. performed the analysis of single-cell RNA sequencing data. J.J.Z. supervised the overall research and experimental execution. Y.N. drafted the manuscript with assistance from Q.W., and J.J.Z. finalized the manuscript.

## Competing interests

S.O. and Q.W. are coinventors of the patent: Methods for enhancing cancer sensitivity to ADAR1 inhibition (UVA LVG Docket No. 182506.00033). Q.L. serves on the Board of Directors of One Patient One Cure and is the owner of Real BioConsulting. J.J.Z. is co-founder and director of Crimson Biopharm Inc. and Geode Therapeutics Inc. All other authors declare no competing interests.

---

**Extended Data Fig. 1.**
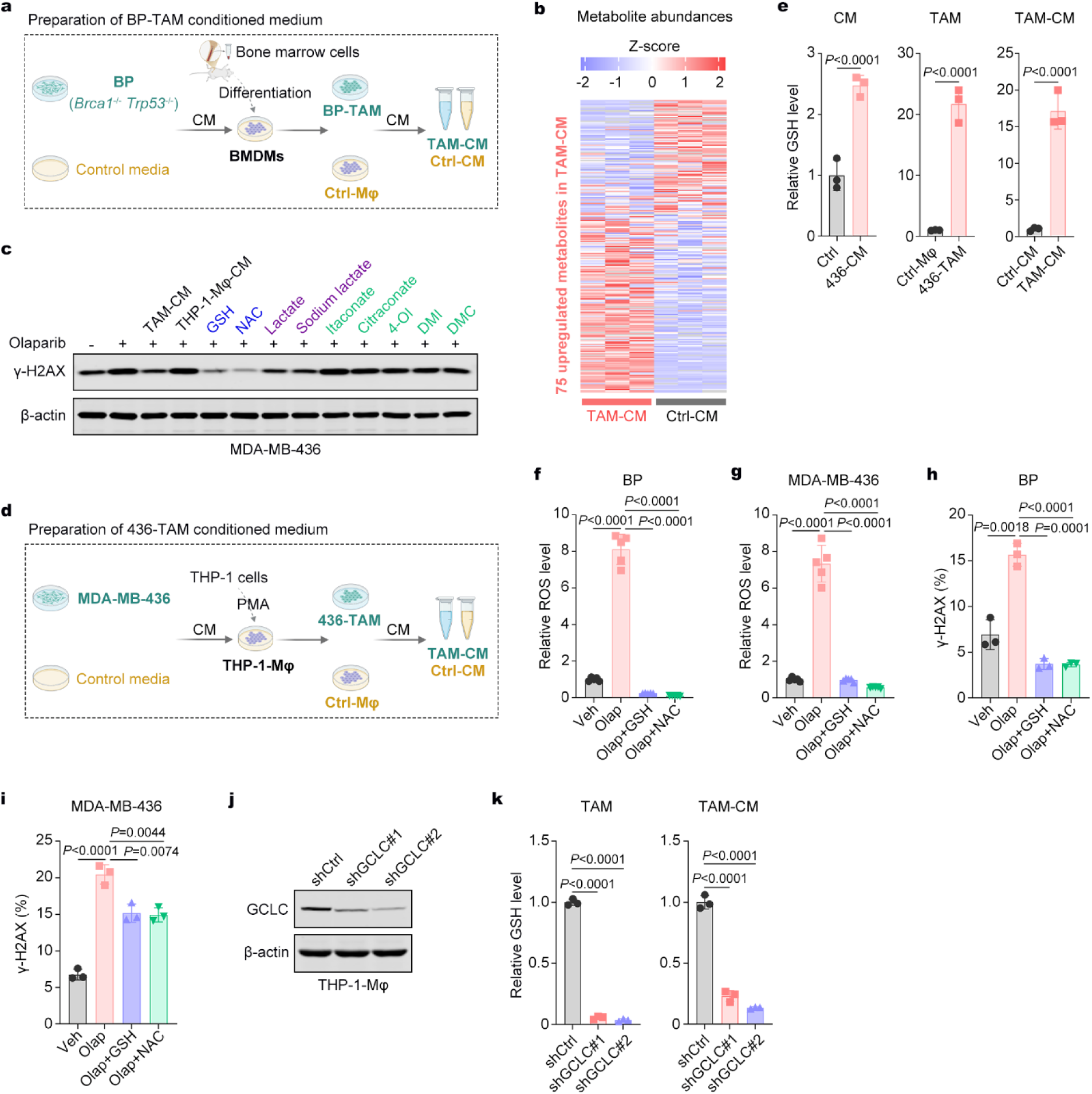
TAM-secreted GSH confers PARPi resistance in BRCA1-deficient breast cancer cells. **a**, Schematic representation of the workflow for the preparation of BP-TAM and CM. BMDMs were incubated with CM from BP cells and control media for 48 hours, and then the CM was collected. **b,** Heatmap illustrates relative metabolite abundances in TAM-CM and Ctrl-CM detected by untargeted metabolomics. **c,** Western blot analysis of γ-H2AX expression in MDA-MB-436 cells following the indicated treatments. Cells were treated with olaparib (20 μM) for 48 hours, and the indicated metabolites, TAM-CM, and THP-1 macrophage-CM were added 12 hours before olaparib treatment. The treatment concentrations were GSH and NAC at 5 mM, lactate and sodium lactate at 20 mM, and itaconate and its derivatives at 200 μM. **d,** Schematic representation of the workflow for the preparation of 436-TAM and CM. THP-1 macrophages were incubated with CM from MDA-MB-436 cells as well as control media for 48 hours and then the CM was collected. **e,** Relative GSH levels in tumor cell-derived CM, TAM and TAM-CM. Data are presented as mean ± s.d.; unpaired Student’s t-test; n = 3. **f,g,** Intracellular ROS levels in BP and MDA-MB-436 cells after olaparib (10 μM) treatment for 48 hours. GSH (5 mM) and NAC (5 mM) were pre-treated for 12 hours before olaparib treatment. Data are presented as mean ± s.d.; unpaired Student’s t-test; n = 5. **h,i,** γ-H2AX expression was detected by flow cytometry in BP and MDA-MB-436 cells after olaparib (10 μM) treatment for 48 hours. GSH (5 mM) and NAC (5 mM) were pre-treated for 12 hours before olaparib treatment. Data are presented as mean ± s.d.; unpaired Student’s t-test; n = 3. **j,** Western blot analysis of GCLC protein expression in THP-1-derived macrophages. **k,** Relative GSH levels in control and GCLC silenced TAM and TAM-CM. Data are presented as mean ± s.d.; unpaired Student’s t-test; n = 3.

**Extended Data Fig. 2.**
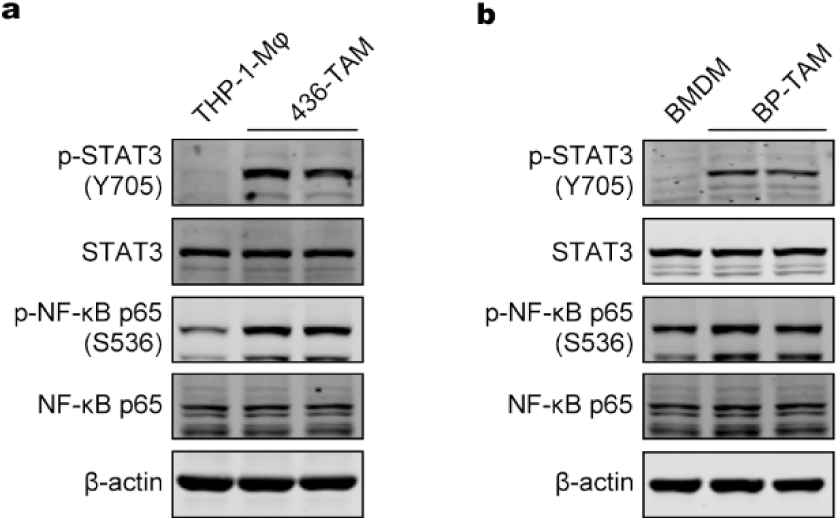
STAT5 signaling activates IRG1 in TAMs. **a,b**, Western blot analysis of STAT3, phosphorylated STAT3 (p-STAT3), NF-κB, and phosphorylated NF-κB (p-NF-κB) expression in THP-1-Mφ, 436-TAMs, BMDMs, and BP-TAMs.

**Extended Data Fig. 3.**
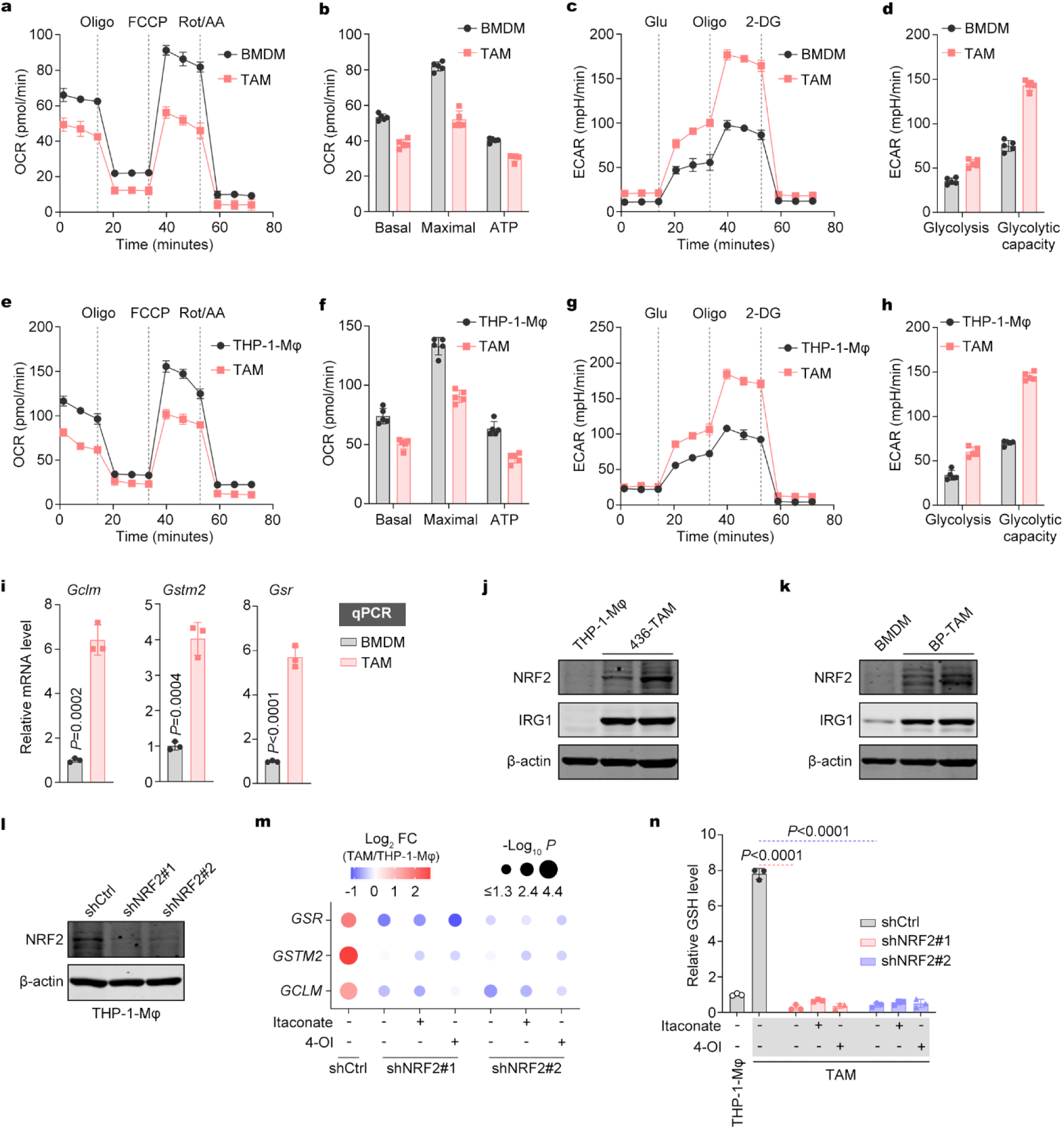
IRG1/itaconate rewires mitochondrial metabolism and promotes GSH synthesis in TAMs. **a,b**, Seahorse assays measuring mitochondrial respiration in BMDMs and BP-TAMs. Data are presented as mean ± s.d.; n = 5. Oligo, oligomycin; FCCP, carbonyl cyanide *p*-trifluoromethoxyphenylhydrazone; Rot/AA, rotenone and antimycin A; OCR, oxygen consumption rate. **c,d,** Seahorse assays detecting glycolysis in BMDMs and BP-TAMs. Data are presented as mean ± s.d.; n = 5. Glu, glucose; Oligo, oligomycin; 2-DG, 2-deoxy-D-glucose; ECAR, extracellular acidification rate. **e,f,** Seahorse assays measuring mitochondrial respiration in THP-1-Mφ and 436-TAMs. Data are presented as mean ± s.d.; n = 5. **g,h,** Seahorse assays detecting glycolysis in THP-1-Mφ and 436-TAMs. Data are presented as mean ± s.d.; n = 5. **i,** RT-qPCR analysis of *Gclm*, *Gstm2*, and *Gsr* mRNA levels in BMDMs and BP-TAMs. Data are presented as mean ± s.d.; unpaired Student’s t-test; n = 3. **j,k,** Western blot analysis of IRG1 and NRF2 protein expression in THP-1-Mφ, 436-TAMs, BMDMs, and BP-TAMs. **l,** Western blot analysis of NRF2 protein expression in control and NRF2-silenced THP-1-Mφ. **m,** RT-qPCR analysis of *GCLM*, *GSTM2*, and *GSR* mRNA expression in control and NRF2-silenced 436-TAMs, with or without itaconate (200 μM) or 4-OI (200 μM) treatment for 12 hours. Fold changes and P values of TAMs relative to control macrophages are shown. **n,** Relative GSH levels in control and NRF2-silenced 436-TAMs, with or without itaconate (200 μM) or 4-OI (200 μM) treatment for 12 hours. Data are presented as mean ± s.d.; unpaired Student’s t-test; n = 3.

**Extended Data Fig. 4.**
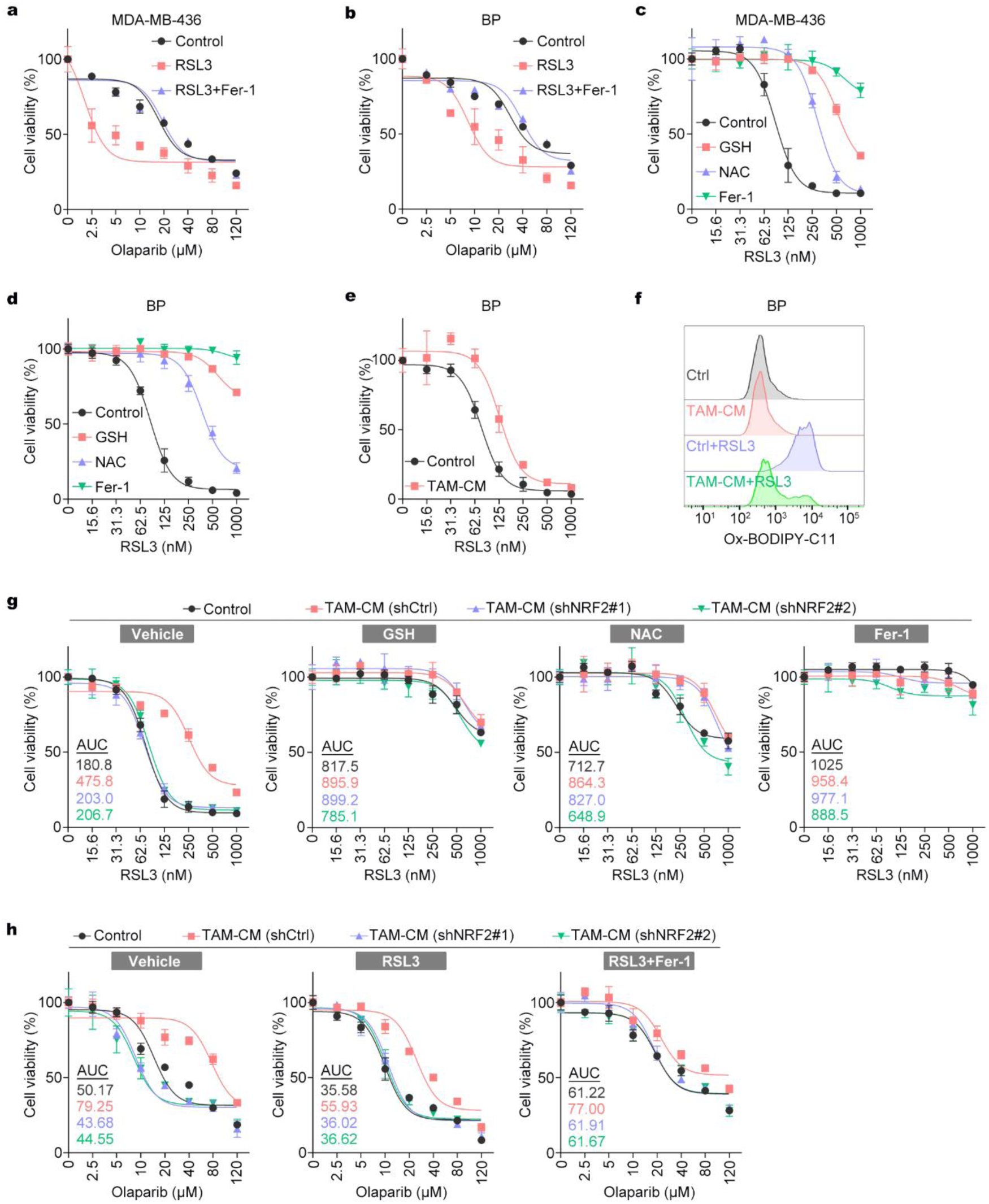
TAM-derived GSH inhibits ferroptosis elicited by combined GPX4 and PARP inhibition. **a,b**, Drug sensitivity assay assessing olaparib-induced cytotoxicity in MDA-MB-436 and BP cells after pretreatment with Fer-1 (10 μM) for 24 hours, followed by co-treatment with RSL3 (100 nM) and olaparib at the indicated concentrations for 48 hours before cell viability was assessed. Data are presented as mean ± s.d.; n = 3. **c,d,** Drug sensitivity assay assessing RSL3-induced cytotoxicity in MDA-MB-436 and BP cells after pretreatment with GSH (5 mM), NAC (5 mM), and Fer-1 (10 μM) for 24 hours, followed by co-treatment with RSL3 at the indicated concentrations for 24 hours before cell viability was assessed. Data are presented as mean ± s.d.; n = 3. **e,** Drug sensitivity assay assessing RSL3-induced cytotoxicity in BP cells after pretreatment with control or TAM-CM for 24 hours. RSL3 was then treated at the indicated concentrations for an additional 24 hours before cell viability was measured. Data are presented as mean ± s.d.; n = 3. **f,** Flow cytometry analysis of lipid peroxidation levels in BP cells pretreated with TAM-CM for 24 hours, followed by co-treatment with RSL3 (250 nM) for 3 hours. **g,** Dose-response curves and AUC values for MDA-MB-436 cells treated with RSL3 at the indicated concentrations for 24 hours. Treatment conditions were as described in Fig. 4 (**d**). Data are presented as mean ± s.d.; n = 3. **h**, Dose-response curves and AUC values for MDA-MB-436 cells treated with olaparib at the indicated concentrations for 48 hours. Treatment conditions were as described in Fig. 4 (**f**). Data are presented as mean ± s.d.; n = 3.

**Extended Data Fig. 5.**
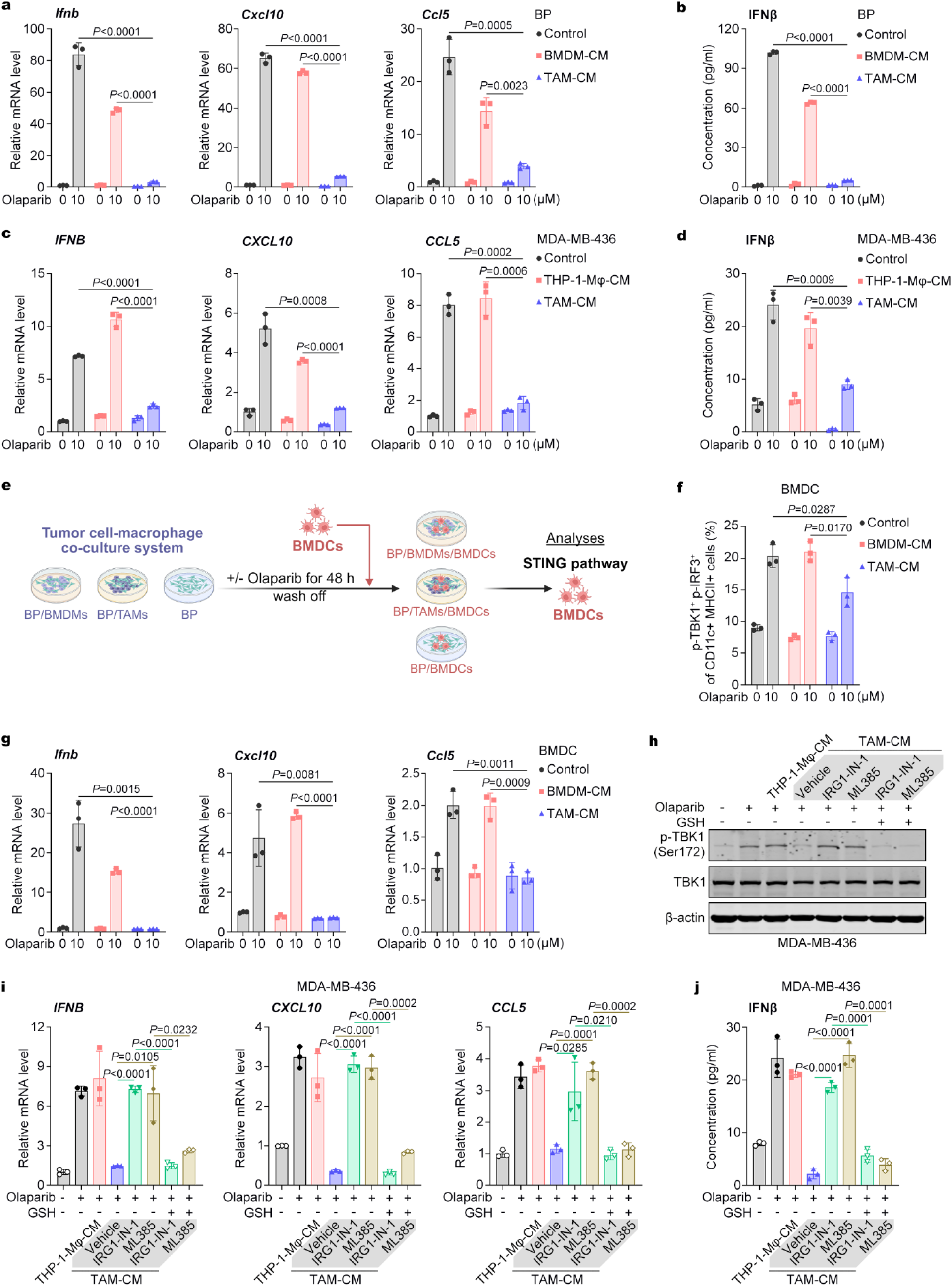
IRG1/NRF2-driven GSH production in TAMs blocks PARPi-induced STING activation in cancer cells and DCs. **a**, RT-qPCR analysis of *Ifnb*, *Cxcl10*, and *Ccl5* mRNA levels in BP cells treated with TAM-CM or BMDM-CM along with olaparib (10 μM) for 48 hours. Data are presented as mean ± s.d.; unpaired Student’s t-test; n = 3. **b,** ELISA quantification of IFNβ levels in BP cell supernatants following olaparib (10 μM) and TAM-CM or BMDM-CM treatment for 48 hours. Data are presented as mean ± s.d.; unpaired Student’s t-test; n = 3. **c,** RT-qPCR analysis of *IFNB*, *CXCL10*, and *CCL5* mRNA levels in MDA-MB-436 cells treated with TAM-CM or THP-1-Mφ-CM along with olaparib (10 μM) for 48 hours. Data are presented as mean ± s.d.; unpaired Student’s t-test; n = 3. **d,** ELISA quantification of IFNβ levels in MDA-MB-436 cell supernatants following olaparib (10 μM) and TAM-CM or THP-1-Mφ-CM treatment for 48 hours. Data are presented as mean ± s.d.; unpaired Student’s t-test; n = 3. **e,** Schematic of the experimental workflow for (**f**) and (**g**). BMDMs or TAMs were co-cultured with BP cells and treated with olaparib (10 μM) for 48 hours. Differentiated BMDCs were then added and incubated for an additional 24 hours before BMDCs were collected for STING pathway analysis. **f,** Flow cytometry analysis of p-TBK1 and p-IRF3-positive BMDCs collected after the experimental workflow described in (**e**). Data are presented as mean ± s.d.; unpaired Student’s t-test; n = 3. **g,** RT-qPCR analysis of *Ifnb*, *Cxcl10*, and *Ccl5* mRNA levels in BMDCs collected after the experimental workflow described in (**e**). Data are presented as mean ± s.d.; unpaired Student’s t-test; n = 3. **h,** Western blot analysis of TBK1 and p-TBK1 expression in MDA-MB-436 cells treated with olaparib (10 μM) and the indicated TAM-CM for 48 hours. GSH (5 mM) was added 4 hours before olaparib treatment. **i,** RT-qPCR analysis of *IFNB*, *CXCL10*, and *CCL5* mRNA levels in MDA-MB-436 cells treated with olaparib (10 μM) and the indicated TAM-CM for 48 hours. GSH (5 mM) was added 4 hours before olaparib treatment. Data are presented as mean ± s.d.; unpaired Student’s t-test; n = 3. **j,** ELISA quantification of IFNβ levels in MDA-MB-436 cell supernatants following olaparib (10 μM) and TAM-CM treatment for 48 hours. GSH (5 mM) was added 4 hours before olaparib treatment. Data are presented as mean ± s.d.; unpaired Student’s t-test; n = 3.

**Extended Data Fig. 6.**
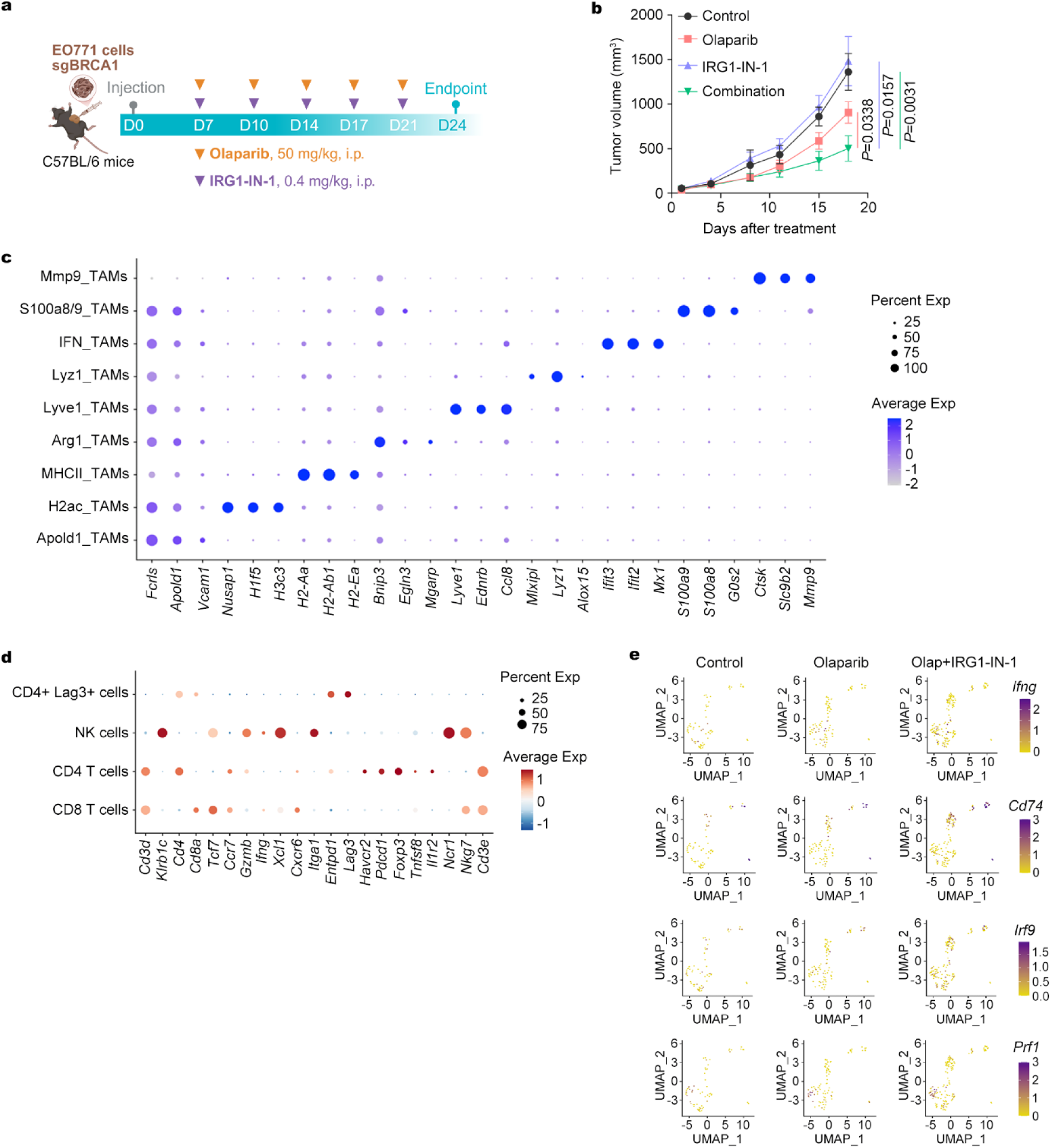
Targeting IRG1 synergizes with PARPi to suppress BRCA1-deficient breast tumors and reshapes TME. **a,b**, Treatment regimen and tumor growth curves of tumors derived from BRCA1-knockout EO771 cells. Treatment was terminated when tumors in any group reached 2000 mm^3^. Data are presented as mean ± s.e.m.; two-way ANOVA; n = 8. **c,** Bubble plot showing relative expression of key marker genes used to define TAM subclusters across treatment groups (vehicle, olaparib alone, or olaparib plus IRG1-IN-1). **d,** Bubble plot displaying relative expression of representative marker genes used to define T/NK cell subclusters. **e,** UMAP projection of T/NK cells showing expression of *Ifng*, *Cd74*, *Irf9*, and *Prf1*.

## References

1. Robson, M. et al. Olaparib for Metastatic Breast Cancer in Patients with a Germline BRCA Mutation. N Engl J Med 377, 523–533 (2017).

2. Tutt, A.N.J. et al. Adjuvant Olaparib for Patients with BRCA1-or BRCA2-Mutated Breast Cancer. N Engl J Med 384, 2394–2405 (2021).

3. Robson, M.E. et al. OlympiAD final overall survival and tolerability results: Olaparib versus chemotherapy treatment of physician’s choice in patients with a germline BRCA mutation and HER2-negative metastatic breast cancer. Ann Oncol 30, 558–566 (2019).

4. Litton, J.K. et al. Talazoparib versus chemotherapy in patients with germline BRCA1/2-mutated HER2-negative advanced breast cancer: final overall survival results from the EMBRACA trial. Ann Oncol 31, 1526–1535 (2020).

5. Ding, L. et al. PARP Inhibition Elicits STING-Dependent Antitumor Immunity in Brca1-Deficient Ovarian Cancer. Cell Rep 25, 2972–2980 e2975 (2018).

6. Wang, Q. et al. STING agonism reprograms tumor-associated macrophages and overcomes resistance to PARP inhibition in BRCA1-deficient models of breast cancer. Nat Commun 13, 3022 (2022).

7. Sen, T. et al. Targeting DNA Damage Response Promotes Antitumor Immunity through STING-Mediated T-cell Activation in Small Cell Lung Cancer. Cancer Discov 9, 646–661 (2019).

8. Shen, J. et al. PARPi Triggers the STING-Dependent Immune Response and Enhances the Therapeutic Efficacy of Immune Checkpoint Blockade Independent of BRCAness. Cancer Res 79, 311–319 (2019).

9. Farkkila, A. et al. Immunogenomic profiling determines responses to combined PARP and PD-1 inhibition in ovarian cancer. Nat Commun 11, 1459 (2020).

10. Long, L.L. et al. PARP Inhibition Induces Synthetic Lethality and Adaptive Immunity in LKB1-Mutant Lung Cancer. Cancer Res 83, 568–581 (2023).

11. Luo, Y. et al. Neoadjuvant PARPi or chemotherapy in ovarian cancer informs targeting effector Treg cells for homologous-recombination-deficient tumors. Cell 187, 4905–4925 e4924 (2024).

12. Wang, S. et al. Gasdermin C sensitizes tumor cells to PARP inhibitor therapy in cancer models. J Clin Invest 134 (2024).

13. Ran, X. et al. PARP inhibitor radiosensitization enhances anti-PD-L1 immunotherapy through stabilizing chemokine mRNA in small cell lung cancer. Nat Commun 16, 2166 (2025).

14. Wagner, J. et al. A Single-Cell Atlas of the Tumor and Immune Ecosystem of Human Breast Cancer. Cell 177, 1330–1345 e1318 (2019).

15. Zhu, Y. et al. Cancer cell-derived arginine fuels polyamine biosynthesis in tumor-associated macrophages to promote immune evasion. Cancer Cell 43, 1045–1060 e1047 (2025).

16. Mehta, A.K. et al. Targeting immunosuppressive macrophages overcomes PARP inhibitor resistance in BRCA1-associated triple-negative breast cancer. Nat Cancer 2, 66–82 (2021).

17. Kloosterman, D.J. & Akkari, L. Macrophages at the interface of the co-evolving cancer ecosystem. Cell 186, 1627–1651 (2023).

18. Ma, R.Y., Black, A. & Qian, B.Z. Macrophage diversity in cancer revisited in the era of single-cell omics. Trends Immunol 43, 546–563 (2022).

19. Nasir, I. et al. Tumor macrophage functional heterogeneity can inform the development of novel cancer therapies. Trends Immunol 44, 971–985 (2023).

20. Yang, M., McKay, D., Pollard, J.W. & Lewis, C.E. Diverse Functions of Macrophages in Different Tumor Microenvironments. Cancer Res 78, 5492–5503 (2018).

21. Klemm, F. et al. Compensatory CSF2-driven macrophage activation promotes adaptive resistance to CSF1R inhibition in breast-to-brain metastasis. Nat Cancer 2, 1086–1101 (2021).

22. Nalio Ramos, R., et al. Tissue-resident FOLR2(+) macrophages associate with CD8(+) T cell infiltration in human breast cancer. Cell 185, 1189–1207 e1125 (2022).

23. Onkar, S. et al. Immune landscape in invasive ductal and lobular breast cancer reveals a divergent macrophage-driven microenvironment. Nat Cancer 4, 516–534 (2023).

24. Pantelidou, C. et al. PARP Inhibitor Efficacy Depends on CD8(+) T-cell Recruitment via Intratumoral STING Pathway Activation in BRCA-Deficient Models of Triple-Negative Breast Cancer. Cancer Discov 9, 722–737 (2019).

25. Ding, L. et al. STING agonism overcomes STAT3-mediated immunosuppression and adaptive resistance to PARP inhibition in ovarian cancer. J Immunother Cancer 11 (2023).

26. Li, X. et al. C5aR1 inhibition reprograms tumor associated macrophages and reverses PARP inhibitor resistance in breast cancer. Nat Commun 15, 4485 (2024).

27. Stenken, J.A. & Poschenrieder, A.J. Bioanalytical chemistry of cytokines--a review. Anal Chim Acta 853, 95–115 (2015).

28. Srinivas, U.S., Tan, B.W.Q., Vellayappan, B.A. & Jeyasekharan, A.D. ROS and the DNA damage response in cancer. Redox Biol 25, 101084 (2019).

29. Peace, C.G. & O’Neill, L.A. The role of itaconate in host defense and inflammation. J Clin Invest 132 (2022).

30. Runtsch, M.C. et al. Itaconate and itaconate derivatives target JAK1 to suppress alternative activation of macrophages. Cell Metab 34, 487–501 e488 (2022).

31. Lin, H. et al. Itaconate transporter SLC13A3 impairs tumor immunity via endowing ferroptosis resistance. Cancer Cell 42, 2032–2044 e2036 (2024).

32. Fan, Y. et al. Itaconate transporter SLC13A3 confers immunotherapy resistance via alkylation-mediated stabilization of PD-L1. Cell Metab (2025).

33. Chen, Y.J. et al. Targeting IRG1 reverses the immunosuppressive function of tumor-associated macrophages and enhances cancer immunotherapy. Sci Adv 9, eadg0654 (2023).

34. Zhao, Y. et al. Neutrophils resist ferroptosis and promote breast cancer metastasis through aconitate decarboxylase 1. Cell Metab 35, 1688–1703 e1610 (2023).

35. Muri, J. & Kopf, M. Redox regulation of immunometabolism. Nat Rev Immunol 21, 363–381 (2021).

36. Mills, E.L. et al. Itaconate is an anti-inflammatory metabolite that activates Nrf2 via alkylation of KEAP1. Nature 556, 113–117 (2018).

37. Lei, G. et al. BRCA1-Mediated Dual Regulation of Ferroptosis Exposes a Vulnerability to GPX4 and PARP Co-Inhibition in BRCA1-Deficient Cancers. Cancer Discov 14, 1476–1495 (2024).

38. Xie, X. et al. Targeting GPX4-mediated ferroptosis protection sensitizes BRCA1-deficient cancer cells to PARP inhibitors. Redox Biol 76, 103350 (2024).

39. Nakamura, T. & Conrad, M. Exploiting ferroptosis vulnerabilities in cancer. Nat Cell Biol 26, 1407–1419 (2024).

40. Mantovani, A., Marchesi, F., Malesci, A., Laghi, L. & Allavena, P. Tumour-associated macrophages as treatment targets in oncology. Nat Rev Clin Oncol 14, 399–416 (2017).

41. Mantovani, A., Allavena, P., Marchesi, F. & Garlanda, C. Macrophages as tools and targets in cancer therapy. Nat Rev Drug Discov 21, 799–820 (2022).

42. Ries, C.H. et al. Targeting tumor-associated macrophages with anti-CSF-1R antibody reveals a strategy for cancer therapy. Cancer Cell 25, 846–859 (2014).

43. Fujiwara, T. et al. CSF1/CSF1R Signaling Inhibitor Pexidartinib (PLX3397) Reprograms Tumor-Associated Macrophages and Stimulates T-cell Infiltration in the Sarcoma Microenvironment. Mol Cancer Ther 20, 1388–1399 (2021).

44. Hamon, P. et al. TGFbeta receptor inhibition unleashes interferon-beta production by tumor-associated macrophages and enhances radiotherapy efficacy. J Immunother Cancer 10 (2022).

45. Xia, Y. et al. TGFbeta reprograms TNF stimulation of macrophages towards a non-canonical pathway driving inflammatory osteoclastogenesis. Nat Commun 13, 3920 (2022).

46. Papadopoulos, K.P. et al. First-in-Human Study of AMG 820, a Monoclonal Anti-Colony-Stimulating Factor 1 Receptor Antibody, in Patients with Advanced Solid Tumors. Clin Cancer Res 23, 5703–5710 (2017).

47. Gomez-Roca, C.A. et al. Phase I study of emactuzumab single agent or in combination with paclitaxel in patients with advanced/metastatic solid tumors reveals depletion of immunosuppressive M2-like macrophages. Ann Oncol 30, 1381–1392 (2019).

48. Kuemmel, S. et al. A Randomized Phase II Study of Anti-CSF1 Monoclonal Antibody Lacnotuzumab (MCS110) Combined with Gemcitabine and Carboplatin in Advanced Triple-Negative Breast Cancer. Clin Cancer Res 28, 106–115 (2022).

49. Rodell, C.B. et al. TLR7/8-agonist-loaded nanoparticles promote the polarization of tumour-associated macrophages to enhance cancer immunotherapy. Nat Biomed Eng 2, 578–588 (2018).

50. Sanlorenzo, M. et al. Systemic IFN-I combined with topical TLR7/8 agonists promotes distant tumor suppression by c-Jun-dependent IL-12 expression in dendritic cells. Nat Cancer 6, 175–193 (2025).

51. Janku, F. et al. Preclinical Characterization and Phase I Study of an Anti-HER2-TLR7 Immune-Stimulator Antibody Conjugate in Patients with HER2+ Malignancies. Cancer Immunol Res 10, 1441–1461 (2022).

52. Dixon, S.J. et al. Ferroptosis: an iron-dependent form of nonapoptotic cell death. Cell 149, 1060–1072 (2012).

53. Lang, X. et al. Radiotherapy and Immunotherapy Promote Tumoral Lipid Oxidation and Ferroptosis via Synergistic Repression of SLC7A11. Cancer Discov 9, 1673–1685 (2019).

54. Wang, W. et al. CD8(+) T cells regulate tumour ferroptosis during cancer immunotherapy. Nature 569, 270–274 (2019).

55. Liao, P. et al. CD8(+) T cells and fatty acids orchestrate tumor ferroptosis and immunity via ACSL4. Cancer Cell 40, 365–378 e366 (2022).

56. Yang, F. et al. Ferroptosis heterogeneity in triple-negative breast cancer reveals an innovative immunotherapy combination strategy. Cell Metab 35, 84–100 e108 (2023).

57. Ran, F.A. et al. Genome engineering using the CRISPR-Cas9 system. Nat Protoc 8, 2281–2308 (2013).

